# EFN-4/Ephrin converges with SAX-3/Robo, UNC-6/Netrin, and Heparan Sulfate Proteoglycan signaling to control MAB-5/Hox-dependent posterior Q neuroblast migration in *Caenorhabditis elegans*

**DOI:** 10.64898/2026.03.27.714887

**Authors:** Vedant D. Jain, Andrew M. Johannesen, Felipe L. Teixeira, Erik A. Lundquist

## Abstract

Hox genes have been broadly implicated in nervous system development, but the molecular and genetic mechanisms that act downstream of Hox factors remain to be identified. The MAB-5 antennapedia-like Hox transcription factor is both necessary and sufficient to cause posterior migration of the Q neuroblast descendants in *Caenorhabditis elegans*. In response to MAB-5, the left-side QL descendants QL.a and QL.ap undergo a three-stage migration process, with each stage characterized by a posterior lamellipodial protrusion followed by cell body migration. The QL.ap cell differentiates into the PQR neuron posterior to the anus. Previous studies showed that the MAB-5-regulated gene *efn-4/Ephrin* was required for the third and final stage of QL.ap migration, with *efn-4* mutation resulting in placement of PQR immediately anterior to the anus. This subtle and previously-undescribed phenotype opens the possibility that other known neuronal development genes could be involved. In this work, we screened known signaling mutants for third-stage PQR migration defects. We found that mutations in SAX-3/Robo signaling, UNC-6/Netrin signaling, and heparan sulfate proteoglycans (HSPGs) all displayed third-stage PQR migration defects. The effects in single mutants were weak compared to *efn-4*, and double mutant analysis revealed lack of genetic synergy, consistent with all of these molecules converging on a common pathway. This genetic analysis is consistent with physical interaction studies *in vitro* from another group that suggest that these molecules form connected communities of interacting extracellular domains, raising the possibility that they are all components of a large extracellular signaling complex required for posterior QL.ap migration. In this model, we envision that MAB-5/Hox drives EFN-4/Ephrin expression in QL.ap, which then seeds the formation of an extracellular signaling complex containing SAX-3/Robo signaling, UNC-6/Netrin signaling, and HSPGs that drives posterior lamellipodial formation and posterior migration.

## Introduction

Neurons and neuroblasts undergo directed migrations in the developing nervous system that are critical to the establishment of circuits and networks. The bilateral Q neuroblasts QL and QR in *Caenorhabditis elegans* are an excellent system in which to define genetic and molecular mechanisms that direct neuroblast and neuronal migration (MIDDELKOOP AND KORSWAGEN 2014; PAOLILLO *et al*. 2024; JAIN AND LUNDQUIST 2025; TEIXEIRA *et al*. 2025). The MAB-5 antennapedia-like Hox transcription factor is both necessary and sufficient for posterior Q neuroblast lineage migration (CHALFIE *et al*. 1983; KENYON 1986; SALSER AND KENYON 1992; HARRIS *et al*. 1996; WHANGBO AND KENYON 1999; KORSWAGEN *et al*. 2000; HERMAN 2003; EISENMANN 2005; JI *et al*. 2013; TAMAYO *et al*. 2013). Hox genes are broadly implicated in nervous system development in vertebrates and invertebrates including *C. elegans* (FENG et al. 2021; SMITH AND KRATSIOS 2024), but the precise molecular and genetic mechanisms employed by Hox genes to control neuronal development remain to be delineated.

The Q cells are born in posterior lateral region in late embryogenesis. Upon hatching, the Q neuroblasts begin division and migration in the L1 larva. Through an identical pattern of division and programmed cell death, each produces three neurons (SULSTON AND HORVITZ 1977; CHALFIE AND SULSTON 1981; MIDDELKOOP AND KORSWAGEN 2014). However, QR and its descendants migrate anteriorly, and QL and descendants migrate posteriorly. In the L1 larva, the Q cells undergo initial protrusion and migration, and by 4-5h after hatching, QL has migrated posteriorly over the V5 seam cell, and QR has migrated anteriorly over the V4 seam cell (HONIGBERG AND KENYON 2000; CHAPMAN *et al*. 2008; MIDDELKOOP AND KORSWAGEN 2014). After this initial migration, the first cell division occurs in the anterior-posterior axis which produces anterior and posterior daughters (QR.a/p and QL.a/p). At this point, QL responds to an EGL-20/Wnt signal from the posterior that, via canonical Wnt signaling, drives expression of the MAB-5 Antennapedia-like Hox transcription factor in QL.a/p. (CHALFIE *et al*. 1983; KENYON 1986; SALSER AND KENYON 1992; HARRIS *et al*. 1996; WHANGBO AND KENYON 1999; KORSWAGEN *et al*. 2000; HERMAN 2003; EISENMANN 2005; JI *et al*. 2013). QR.a/p do not respond to the EGL-20 signal and do not activate MAB-5 expression. MAB-5 is required for the further posterior migration of QL descendants. After this initial directional decision, Wnts control QL and QR descendant migrations (ZINOVYEVA *et al*. 2008; JOSEPHSON *et al*. 2016; PAOLILLO *et al*. 2024).

AQR (QR.ap) and PQR (QL.ap) undergo the longest-range migrations, with AQR in the anterior deirid near the posterior pharyngeal bulb and PQR in the phasmid ganglia posterior to the anus (WHITE *et al*. 1986; CHAPMAN *et al*. 2008). *egl-20* and *mab-5* loss-of-function mutants have no effect on initial QL and QR protrusion and migration, but QL descendants fail to migrate posteriorly and instead migrate anteriorly (CHAPMAN *et al*. 2008; TAMAYO *et al*. 2013; JOSEPHSON *et al*. 2016; JAIN AND LUNDQUIST 2025). The opposite happens in *mab-5* gain-of-function, where QR and QL descendants migrate to the posterior despite normal initial migration of QR (SALSER AND KENYON 1992; CHAPMAN *et al*. 2008; TAMAYO *et al*. 2013). These studies demonstrate that MAB-5 is a determinant of posterior migration of Q descendants.

To identify genes regulated by MAB-5 in the Q cells, fluorescence-activated Q cell sorting combined with RNA-seq was conducted on *wild-type, mab-5* loss-of-function, and *mab-5* gain-of -function (PAOLILLO *et al*. 2024). Three genes, *vab-8/KIF26, lin-17/Frizzled,* and *efn-4/Ephrin*, were positively regulated by MAB-5 and controlled posterior PQR migration downstream of MAB-5 (PAOLILLO *et al*. 2024; JAIN AND LUNDQUIST 2025). These studies also found that posterior PQR migration involves three distinct stages, each including extension of a posterior lamellipodial protrusion and later migration of the cell body. VAB-8/KIF26 was required for all three stages, LIN-17/Frizzled for stages two and three, and EFN-4/Ephrin for the third and final stage only (JAIN AND LUNDQUIST 2025). MAB-20/Semaphorin, not regulated by MAB-5, acted with EFN-4 to control the third stage (JAIN AND LUNDQUIST 2025). This third-stage migration failure represented a novel and previously-undescribed phenotype where PQR is positioned immediately anterior to the anus.

In this work, we survey known signaling molecules that control nervous system development for the subtle and novel third stage PQR migration defect that might have been overlooked when the mutants were previously analyzed. Indeed, we found that components of SAX-3/Robo signaling, UNC-6/Netrin signaling, and heparan sulfate proteoglycans (HSPGs) were involved. The many mutants displayed only moderate

PQR migration defects on their own, and often did not enhance each other or *efn-4* mutants. This suggests that these molecules might all converge on a common pathway. Furthermore, *efn-4* mutants showed the strongest effect that no other single or double mutant significantly exceeded. A previous *in vitro* Extracellular Interactome Assay (EICA) involving extracellular domain interactions (NAWROCKA *et al*. 2024) showed that these molecules form “connected communities” of interactions, including EFN-4, SAX-3/Robo, SLT-1/Slit, and MAB-20/Semaphorin. Here, AlphaFold3 (VARADI *et al*. 2024) was used to model these potential physical interactions and the possible formation of a multimeric complex including these molecules. Taken together, these data suggest a model in which MAB-5/Hox drives the expression of EFN-4/Ephrin in QL.ap, which seeds the formation of a large extracellular signaling complex that drives posterior lamellipodial protrusion and the third and final migration of QL.ap.

## Results

### The step-wise posterior migration of QL.a and QL.ap defines a new PQR migration phenotype

We showed previously that the posterior migration of QL.a and QL.ap occurs in a three-step process. After QL migrates posteriorly over V5, it undergoes its first division to form QL.a and QL.p (Figure 1A and B). QL.a first protrudes and migrates posteriorly past QL.p, which does not migrate (Figure 1C). QL.a then divides to form QL.aa, which undergoes programmed cell death, and QL.ap, which protrudes and migrates posteriorly a second time to reside in a position immediately anterior to the anus (Figure 1C). QL.ap protrudes and migrates posteriorly a third time over the anus to reside in a position posterior to the anus, where it begins differentiation into the PQR neuron by extending a posterior dendritic process and a ventral axon (Figure 2A). Mutations that affect posterior QL.a migration can have effects at each independent step (JAIN AND LUNDQUIST 2025) (Table 1). *vab-8/KIF26A* mutants affect each step. PQR misplacement from near the place of birth and posterior to QL.a is defined as paradigm 1 PQR migration failures of the first and second migrations. PQR misplacement immediately posterior to the anus was defined as paradigm 2 PQR migration failure at the third and final step (Figure 1C). *lin-17/Frizzled* mutations affected the second and third steps and also showed paradigm 1 and 2 defects. *efn-4/Ephrin* and *mab-20/Semaphorin* specifically affected the third migration and displayed only paradigm 2 defects, with PQR immediately anterior to the anus. None affected AQR migration. These PQR migration defects, particularly the distinctive paradigm 2 defect of *efn-4* and *mab-20*, are previously unappreciated phenotypes and represent an opportunity to discover new genes and pathways that control these migrations.

**Figure 1.**
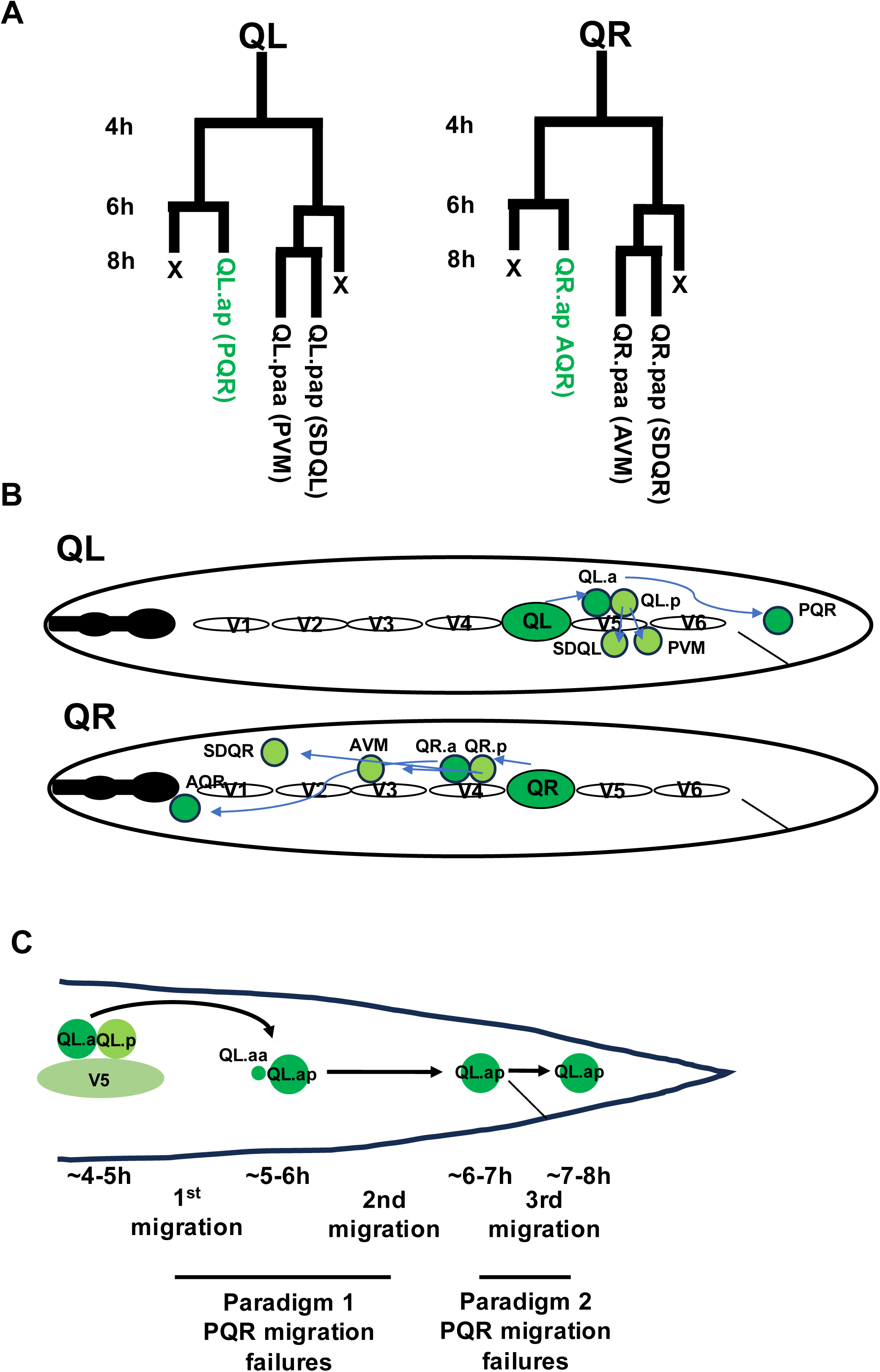
Q lineage migration. A) The lineage map for the QL and QR neuroblast lineages. Post-hatch time is indicated on the left. B) A schematic of migration of the left and right lineage (QL and QR) is shown. In the first phase of migration, QL and QR migrate over the V5 and V4 seam cells respectively. In the second phase of migration, both QL and QR undergo the first round of cell division and begin migration. QL gives rise to the PVM and SDQL neurons which reside near the plase of birth, and to PQR, which migrates posteriorly into the tail behind the anus. QR descendants migrate anteriorly, giving rise to AVM, SDQL, and AQR, which migrates furthest anteriorly to reside in the anterior deirid ganglion near the pharynx. C) Shown is a schematic representing the three distinct phases of QL.a and QL.ap posterior migration as previously described in Jain and Lundquist, 2025. In the first migration, QL.a protrudes and migrates posteriorly over QL.p. QL.a divides, giving rise to QL.aa, which undergoes programmed cell death, and QL.ap, which undergoes the second migration to a position just anterior to the anus. In the third migration, QL.ap protrudes and migrates posteriorly over the anus. In this position posterior to the anus, QL.ap begins differentiation into the PQR neuron. The approximate time after hatching when each migration occurs is indicated. In mutants affecting posterior QL> and QL.apo migration, PQR differentiates at locations of migration failure, which are defined as paradigm I (for failures of the first and second migrations), and paradigm 2, failure of the third migration, resulting in a characteristic placement of the PQR neuron immediately anterior to the anus.

**Figure 2.**
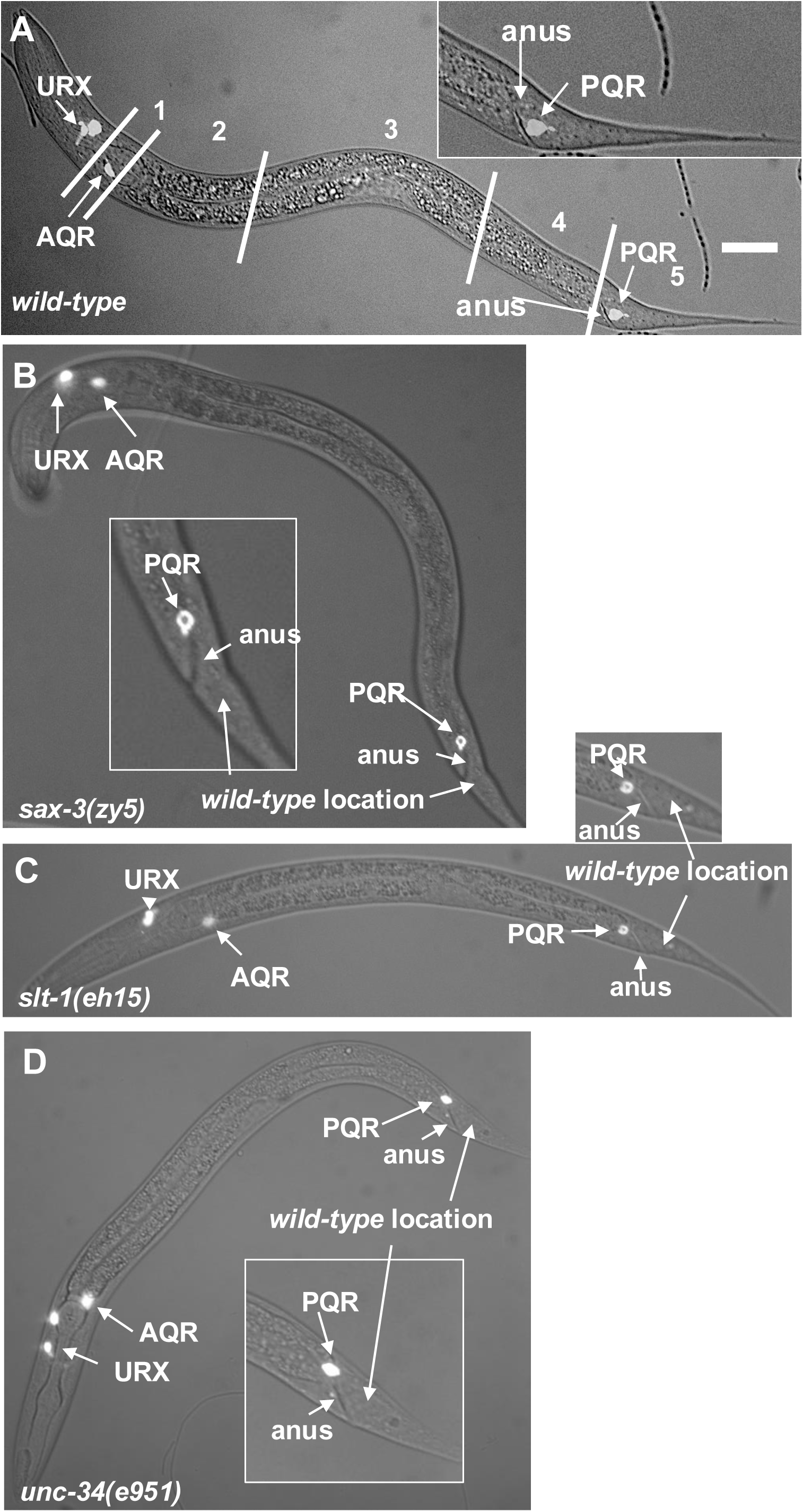
PQR migration defects in *sax-3, slt-1,* and *unc-34* mutants. Fluorescence and differential interference contrast micrographs of L4 larval stage animals with *Pgcy-23::cfp* expression in AQR, PQR, and URX left and right. The URX left and right cell bodies are close together and usually not distinguishable in these micrographs and so their position is indicated by URX. The scale bar in A represents 10µm. The insets are magnified posterior regions of the animals showing the position of PQR relative to the anus. A) A *wild-type* animal. The five regions along the anterior-posterior axis as shown in Table 1 are indicated. The *wild-type* positions of AQR and PQR are positions 1 and 5, respectively. B) A *sax-3(zy5)* mutant animal with PQR mispositioned immediately anterior to the anus (paradigm I PQR migration defect), indicative of a failure in the third and final QL.ap migration. C-D) A *slt-1(eh15)* and *unc-34(e951)* mutant animals with paradigm I PQR migration defects.

**Table 1.** PQR migration defects.

| Genotype<sup>1</sup> | PQR position<sup>2</sup> |  |  |  |  |  |
| --- | --- | --- | --- | --- | --- | --- |
|  | 1 | 2 | 3 | 4.1<br>3 | 4.2<sup>4</sup> | 5 |
| *wild-type* | 0 | 0 | 0 | 0 | 0 | 100 |
| *efn-4(bx80)* | 0 | 0 | 1 | 0 | 70 | 29 |
| *vab-8(ev411)* | 1 | 0 | 1 | 30 | 22 | 48 |
| *lin-17(n671)* | 10 | 4 | 8 | 20 | 28 | 30 |
| *mab-20(ev574)* | 0 | 0 | 0 | 0 | 50 | 50 |
| *lqls320[mab-5scCRISPR]* | - | - | - | 22 | 12 | 66 |
| *lad-2(tm3056)* | 0 | 0 | 0 | 0 | 7 | 93 |
| *lad-2(hd31)* | 0 | 0 | 0 | 0 | 2 | 98 |
| *sax-7(ky146)* | 0 | 0 | 0 | 0 | 7 | 93 |
| *plx-2(ev773)* | 0 | 0 | 0 | 0 | 0 | 100 |
| *plx-2(ev775)* | 0 | 0 | 0 | 0 | 0 | 100 |
| *plx-2(ev773); lad-2(tm3056)* | 0 | 0 | 0 | 0 | 6 | 94 |
| *sax-3(gk5367)* | 0 | 0 | 0 | 10 | 26 | 64 |
| *sax-3(zy5)* | 0 | 0 | 0 | 4 | 14 | 82* |
| *sax-3(ky123)* | 0 | 0 | 0 | 0 | 29 | 71 |
| *sax-3(zy152)* | 0 | 0 | 0 | 10 | 17 | 73 |
| *sax-3(yad175)* | 0 | 0 | 0 | 8 | 55* | 54 |
| *lqls461: Sc-CRISPR for sax-3* | 0 | 0 | 0 | 0 | 9 | 91 |
| *lqls462: Sc-CRISPR for sax-3* | 0 | 0 | 0 | 0 | 10 | 90 |
| *slt-1(eh15)* | 0 | 0 | 0 | 0 | 20 | 80 |
| *slt-1(ev740)* | 0 | 0 | 0 | 0 | 7 | 93 |
| *slt-1(ev741)* | 0 | 0 | 0 | 0 | 2 | 98 |
| *slt-1(gk952391)* | 0 | 0 | 0 | 0 | 11 | 89 |
| *slt-1(gk301595)* | 0 | 0 | 0 | 0 | 13 | 87 |
| *slt-1(tm307)* | 0 | 0 | 0 | 0 | 4 | 96 |
| *sax-3(gk5367); slt-1(eh15)* | 0 | 0 | 0 | 5 | 35 | 60 |
| *sax-3(ky123); slt-1(eh15)* | 0 | 0 | 0 | 3 | 25 | 72 |
| *sax-3(zy152); slt-1(eh15)* | 0 | 0 | 0 | 3 | 17 | 80 |
| *unc-34(e951)* | 0 | 0 | 0 | 15 | 46 | 39 |
| *unc-34(gm104)* | 0 | 0 | 0 | 3 | 47 | 50 |
| *unc-34(lq17)* | 0 | 0 | 0 | 0 | 32 | 68 |
| *unc-34(e951); slt-1(eh15)* | 0 | 0 | 0 | 12 | 64 | 24 |
| *unc-34(e951); sax-3(gk5367)* | 0 | 0 | 0 | 6 | 38* | 56 |
| *unc-34(e951); efn-4(bx80)* | 0 | 0 | 0 | 5 | 65 | 30 |
| *sax-3(gk5367)M+; efn-4(bx80)* | 0 | 0 | 0 | 10 | 53* | 37 |
| *sax-3(zy5); efn-4(bx80)* | 0 | 0 | 0 | 13* | 73 | 17 |
| *sax-3(ky123); efn-4(bx80)* | 0 | 0 | 0 | 10* | 65 | 25 |
| *slt-1(eh15)M+; efn-4(bx80)* | 0 | 0 | 0 | 0 | 82 | 18 |
| *sax-3(zy5)M+; mab-20(ev574)* | 0 | 0 | 0 | 11 | 51 | 38 |
| *slt-1(eh15); mab-20(ev574)M+* | 0 | 0 | 0 | 0 | 69 | 31 |
| *sax-3(gk5367); vab-8(ev411)* | 0 | 0 | 0 | 46 | 34* | 20* |
| *sax-3(zy5); vab-8(ev411)* | 0 | 0 | 0 | 57* | 30* | 13* |
| *sax-3(ky123); vab-8(ev411)* | 0 | 0 | 0 | 45* | 37 | 19* |
| *sax-3(gk5367)M+; lin-17(n671)* | 12 | 6 | 6 | 10 | 43 | 23 |
| *sax-3(zy5); lin-17(n671)* | 8 | 3 | 4 | 37* | 23 | 25 |
| *sax-3(ky123); lin-17(n671)* | 0 | 0 | 0 | 6 | 36 | 58 |
| *unc-6(ev400)* | 0 | 0 | 0 | 10 | 17 | 73 |
| *unc-6(e78)* | 0 | 0 | 0 | 22 | 28 | 50* |
| *unc-40(n324)* | 54 | 1 | 2 | 6 | 11 | 26 |
| *unc-40(e1430)* | 22* | 0 | 0 | 4 | 19 | 55 |
| *unc-5(e53)* | 0 | 0 | 0 | 4 | 22 | 74 |
| *unc-5(ev480)* | 0 | 0 | 0 | 0 | *9 | 91 |
| *unc-5(wy1996; wy1796)* | 0 | 0 | 0 | 0 | 14 | 86 |
| *unc-5(wy1949)* | 0 | 0 | 0 | 0 | 11 | 89 |
| *unc-40(n324); unc-6(ev400)* | 44 | 5 | 7 | 7 | 18 | 19 |
| *unc-40(n324); unc-5(e53)* | 45 | 6 | 6 | 7 | 14 | 22 |
| *unc-6(ev400); sax-3(zy5)M+* | 2 | 2 | 2 | 13 | 26 | 55 |
| *unc-40(n324); sax-3(zy5)M+* | 64 | 2 | 3 | 7 | 17* | 7* |
| *unc-6(ev400); slt-1(eh15)* | 0 | 0 | 0 | 15 | 35 | 50 |
| *unc-40(n324); slt-1(eh15)* | 92* | 0 | 0 | 0 | 3 | 5 |
| *unc-40(n324); slt-1(eh15)* | - | - | - | 24* | 43 | 33* |
| *unc-6(ev400)M+; efn-4(bx80)* | 4 | 0 | 8 | 30* | 33 | 25 |
| *unc-6(e78)M+; efn-4(bx80)* | 0 | 0 | 5 | 21 | 67 | 7 |
| *unc-40(n324)M+; efn-4(bx80)* | 52 | 7* | 17* | 6 | 13 | 5* |
| *unc-40(e1430)M+; efn-4(bx80)* | 33 | 8 | 13 | 13 | 22 | 11 |
| *unc-6(e78)M+; mab-20(ev574)* | 9 | 6 | 7 | 14 | 56 | 8 |

<sup>1</sup>For each genotype, 100 animals were scored on three separate occasions for a total of 300 animals. Except for *unc-40* single and double mutants, no AQR migration defects were observed in these genotypes.
<sup>2</sup>Numbers are average percentages from three replicates of cells at each position rounded to the nearest whole number.
<sup>3</sup>Position 4, paradigm 1 as described in Results.
<sup>4</sup>Position 4, paradigm 2 as described in Results.
\* $p \leq 0.05$ (see Results for description).

The MAB-5/Hox transcription factor is both necessary and sufficient for posterior migration of Q descendants (HARRIS *et al*. 1996; TAMAYO *et al*. 2013; JOSEPHSON *et al*. 2016). Previous work showed that *efn-4, vab-8,* and *lin-17* are MAB-5-regulated genes that act downstream of MAB-5 in posterior migration (PAOLILLO *et al*. 2024; JAIN AND LUNDQUIST 2025). The ability to analyze the role of *mab-5* in posterior PQR migration is masked by the high-penetrance anterior PQR migration defect in existing *mab-5* mutants (JOSEPHSON *et al*. 2016). However, cell-specific Cas9 genome editing of *mab-5* resulted in a lower penetrance of anterior PQR migration (OCHS *et al*. 2020), allowing for potential defects in posterior migration to be scored. Indeed, of the PQR that migrated posteriorly in *lqIs320[mab-5scCRISPR]* animals, 22% paradigm 1 and 12% paradigm 2 PQR migration defects were detected (Table 1). This result confirms that MAB-5 is required to both inhibit anterior QL descendant migration and to promote posterior QL.a and QL.ap migration as previously described (JOSEPHSON *et al*. 2016).

### SAX-3/Robo controls posterior PQR migration

In our previous study, *efn-4* was shown to promote the third and final step of posterior migration of QL.ap downstream of *mab-5* (JAIN AND LUNDQUIST 2025). The sole Ephrin receptor tyrosine kinase VAB-1 was not involved, suggesting a distinct mechanism (JAIN AND LUNDQUIST 2025). EFN-4 interacts genetically with the PLX-2/Plexin receptor (HAHN AND EMMONS 2003; NAKAO *et al*. 2007; IKEGAMI *et al*. 2012). *plx-2* mutants had no PQR migration defects (Table 1), suggesting it is not involved in posterior PQR migration. EFN-4 is known to interact with the Ig-domain-containing receptors LAD-2 and SAX-7 in axon guidance and to interact physically with LAD-2 (DONG *et al*. 2016). *lad-2* and *sax-7* mutants had weak paradigm 2 PQR migration defects (2-7%), suggesting that might be involved in posterior QL.ap migration (Table 1). The role of *lad-2* and *sax-7* in posterior PQR migration will be the subject of future studies.

EFN-4 and the Ig receptor SAX-3 have redundant functions in embryonic morphogenesis, with double mutants displaying an embryonic lethal phenotype (GHENEA *et al*. 2005; MILLER AND CHIN-SANG 2012). Recently, SAX-3/Robo was shown to physically interact with EFN-4 in a high-throughput *in vitro* Extracellular Interactome Assay (ECIA) screen (NAWROCKA et al. 2024). The D2 domain of EFN-4 specifically interacts with the first four extracellular Immunoglobulin domains of SAX-3 (NAWROCKA et al. 2024). Furthermore, our previous studies showed that *sax-3(ky123)* mutants had PQR posterior migration defects (JOSEPHSON *et al*. 2017).

Five *sax-3* mutants were screened for paradigm 1 and 2 PQR defects in the context of the three-step migration pattern. The five alleles, *gk5367, ky123, zy5, zy152, and yad175*, caused paradigm 1 defects (0-10%) and/or paradigm 2 defects (14-55%) (Table 1; Figure 2B). None displayed anterior PQR migration or obvious defects in AQR migration. *sax-3(gk5367)*, a deletion of all the exons except the first and a likely null mutant (AU *et al*. 2019), displayed a combined penetrance of 36%. *sax-3(zy152),* which deletes regions of final exons 9 through 12, showed a combined penetrance of 27%, not significantly different from *sax-3(gk5367)*. *sax-3(zy5)* introduces a premature stop codon in exon 9 (SHAH *et al*. 2017) and showed a combined penetrance of 18%, significantly weaker than *sax-3(gk5367)* (*p* = 0.0065). Possibly, some activity remains in *sax-3(zy5)* due to stop codon read-through, splicing, or incomplete nonsense-mediated decay. *sax-3(ky123)* is a deletion of the first exon (ZALLEN *et al*. 1998) and showed only paradigm 2 defects (29%), similar to *efn-4*. Recently, a novel exon in the large intron upstream of exon 6 was described (WEINREB *et al*. 2025). The exon encodes a signal peptide and is spliced in frame to exon 6. This transcript can encode a truncated SAX-3 molecule lacking Ig domains 1-4, but containing the remainder of the molecule. *sax-3(ky123)* might only affect the long isoform containing Ig domains 1-4, the region that physically interacts with EFN-4. Possibly, the long isoform is required only for the third and final QL.ap migration similar to *efn-4,* whereas the short isoform might be involved in the first, second, and/or third migrations. Indeed, Ig domains 1-4, missing in this isoform, interact physically with EFN-4 (NAWROCKA *et al*. 2024). In any case, this result suggests a long-isoform-specific function of SAX-3.

*sax-3(yad175)* allele, which is an in-frame deletion of the second fibronectin domain of SAX-3, also removes a protease cleavage site that can release a soluble, secreted form of the Ig domains and the first fibronectin domain (QU *et al*. 2020). *sax-3(yad175)* showed significantly increased paradigm 1 defects compared to the *sax-3(gk5367)* null. Possibly, *sax-3(yad175)* is not a simple loss-of-function mutation.

In sum, these results show that SAX-3 is required for multiple stages of posterior PQR migration, with a possible long-isoform-specific role in the third migration. This is consistent with the physical interaction of EFN-4 with the first four Ig domains of SAX-3 present only in the long isoform (NAWROCKA *et al*. 2024).

### *sax-3* can act cell-autonomously in PQR migration

*vab-8, lin-17,* and *efn-4* act downstream of the MAB-5/Hox gene to control QL.a and QL.ap migration (JAIN AND LUNDQUIST 2025). Further analysis of publicly-available single-cell RNA sequencing data of the Q lineage (TEIXEIRA *et al*. 2025) revealed that *efn-4* and *vab-8* were enriched in the QL.a lineage (TEIXEIRA *et al*. 2025), whereas *lin-17* was expressed in both QL.a and QL.p (Figure 3). *mab-20* was not regulated by MAB-5 (PAOLILLO *et al*. 2024) and was also expressed in both QL.a and QL.p. lineages (Figure 3).

**Figure 3:**
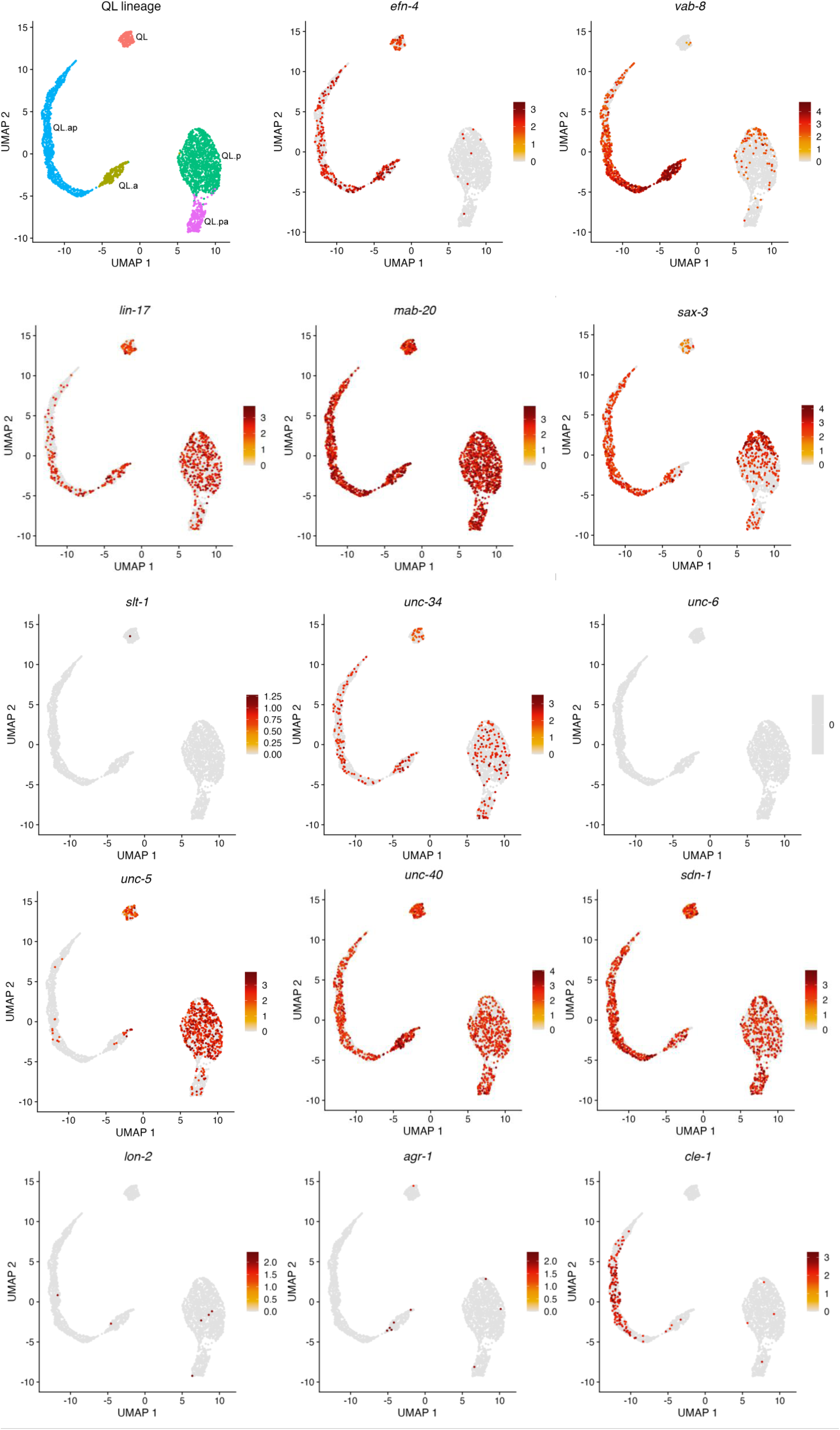
UMAP projections of single-cell RNA sequencing data showing gene expression patterns across the QL lineage. The top-left panel shows annotated QL lineage subpopulations. Each remaining panel displays the expression of an individual gene projected onto the same UMAP coordinates. Cells are colored by expression level (yellow to red scale), with gray indicating undetectable expression.

*sax-3* expression is not regulated by *mab-5* in the Q lineages, but *sax-3* expression is enriched in the Q lineages (PAOLILLO *et al*. 2024). *sax-3* was expressed in both the QL.a and QL.p lineages (Figure 3). To test the autonomy of function of *sax-3* in posterior PQR migration, we used cell-specific Cas9 genome editing as previously described (OCHS *et al*. 2020). Two separate synthetic guide RNAs were made: one for 9^th^ exon and another for 12^th^ exon. These sgRNAs target both long and short isoforms. Each synthetic guide RNA was driven under the U6 promoter, and Cas9 expression was driven under the *egl-17* promoter expressed in the Q lineages. Two independent transgenes, *lqIs461* and *lqIs462*, were scored for migration defects. As shown in Table 2, both transgenes showed a significant migration paradigm 2 PQR posterior migration defect (9% and 10%, respectively). No paradigm 1 defects were observed. This result suggests that *sax-3* can act cell-autonomously in the Q lineages. The low-penetrance phenotype could be due to inefficient genome editing (OCHS *et al*. 2020). Alternatively, *sax-3* might also have non-autonomous effects on PQR migration.

**Table 2.**
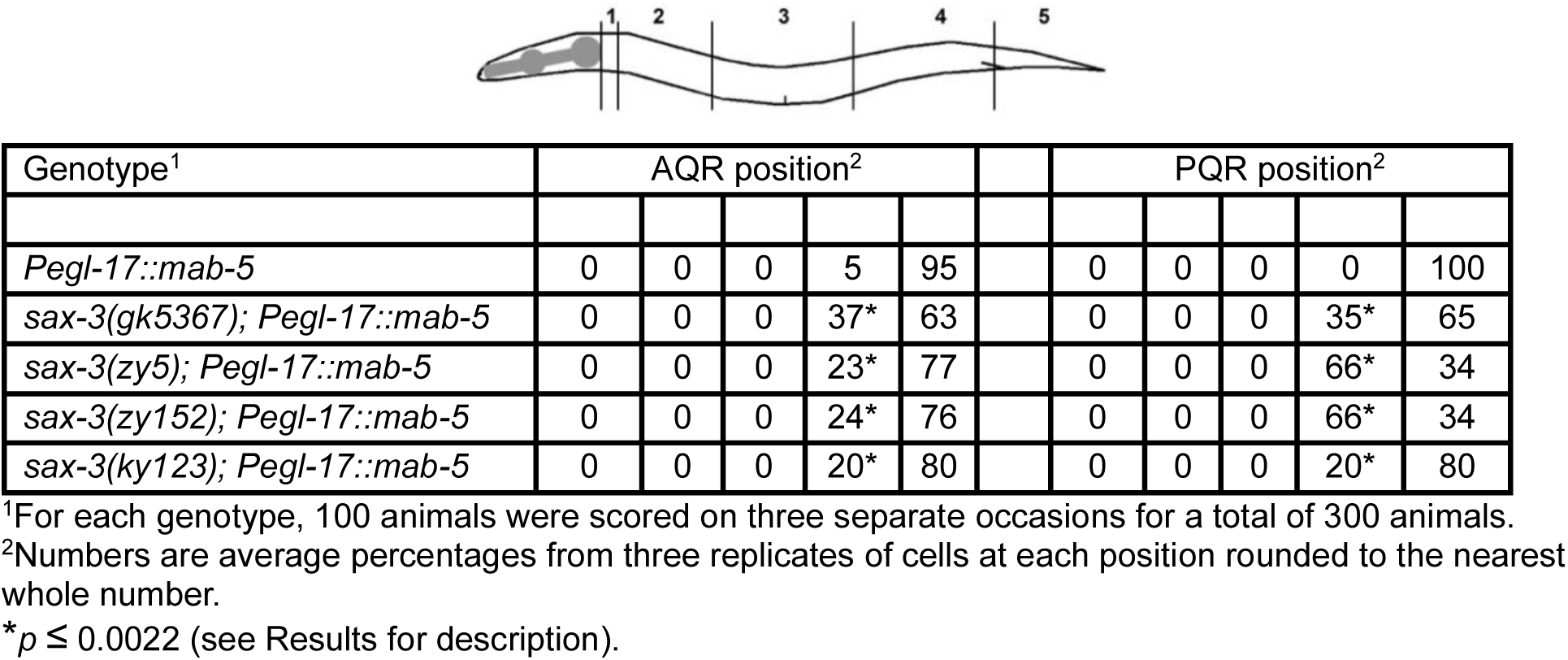
Suppression of MAB-5 gain-of-function in AQR and PQR migration.

### *sax-3* acts downstream of *mab-5*

We wanted to test if *sax-3* is part of the same pathway downstream of *mab-5* as *efn-4, mab-20* and *vab-8* (JAIN AND LUNDQUIST 2025). While *mab-20* is not regulated by *mab-5* (PAOLILLO *et al*. 2024), it is required for the full effect of *mab-5* gain-of-function (JAIN AND LUNDQUIST 2025). The *lqIs221[Pegl-15::mab-5]* transgene drives ectopic *mab-5* expression in both QL and QR lineages, and both AQR and PQR migrate posteriorly to the normal position of PQR (Table 2; Figure 4A) (TAMAYO *et al*. 2013). No AQR migrated anteriorly, and only 5% failed to migrate posterior to the anus as previously described (Table 2). Double mutants of *lqIs221[Pegl-17::mab-5]* and three *sax-3(gk5367, zy5, zy152,* and *ky123)* were scored for AQR and PQR migration (Table 2). Significant suppression of posterior AQR migration was observed, with 20-37% of AQRs failing to migrate posterior to the anus (*p* ≤ 0.0022), although none migrated anteriorly. PQR posterior migration was also affected, with 20-66% of PQRs failing to migrate posterior to the anus, although none migrated anteriorly (Table 2). In two cases, with *sax-3(zy5* and *zy152)* the percentage of PQR is position 4 was significantly higher than in the *sax-3* mutant alone (*p* < 0.0001). We do not understand the nature of this interaction, but it is possible that in some contexts, *mab-5* overexpression can inhibit migration as observed previously (JAIN AND LUNDQUIST 2025). These data indicate that SAX-3 is required for the full effect of MAB-5 in posterior migration.

**Figure 4:**
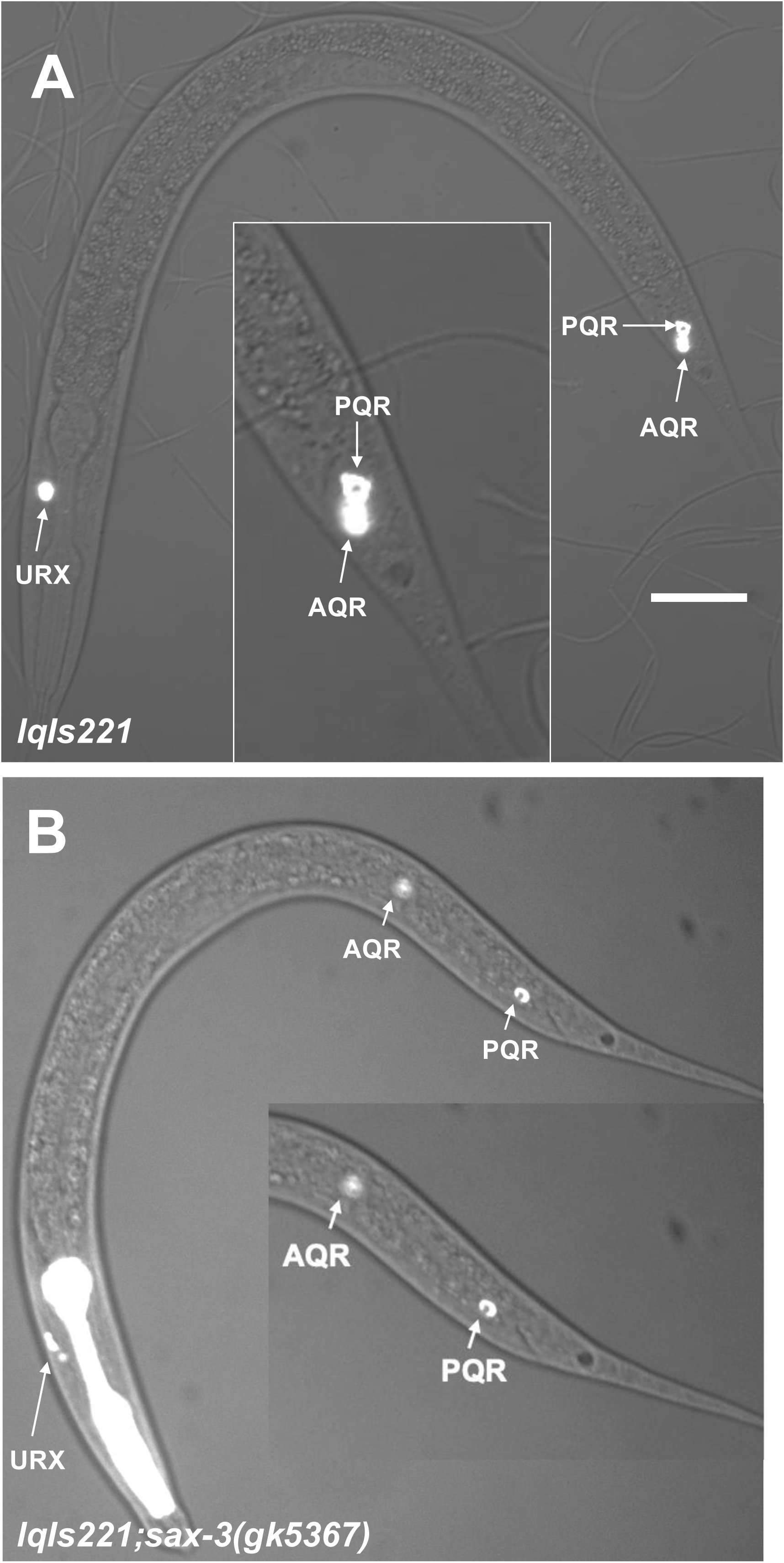
**Suppression of MAB-5 gain-of-function by *sax-3.*** Micrographs as described in Figure 2 are shown. The scale bar in A represents 10µm. A) An *lqIs221[Pegl-17::mab-5]* gain-of-function animal displays both AQR and PQR behind the anus. B) In this *sax-3(gk5367); lqIs221* double mutant, both AQR and PQR are positioned anterior to the anus.

**Figure 5.**
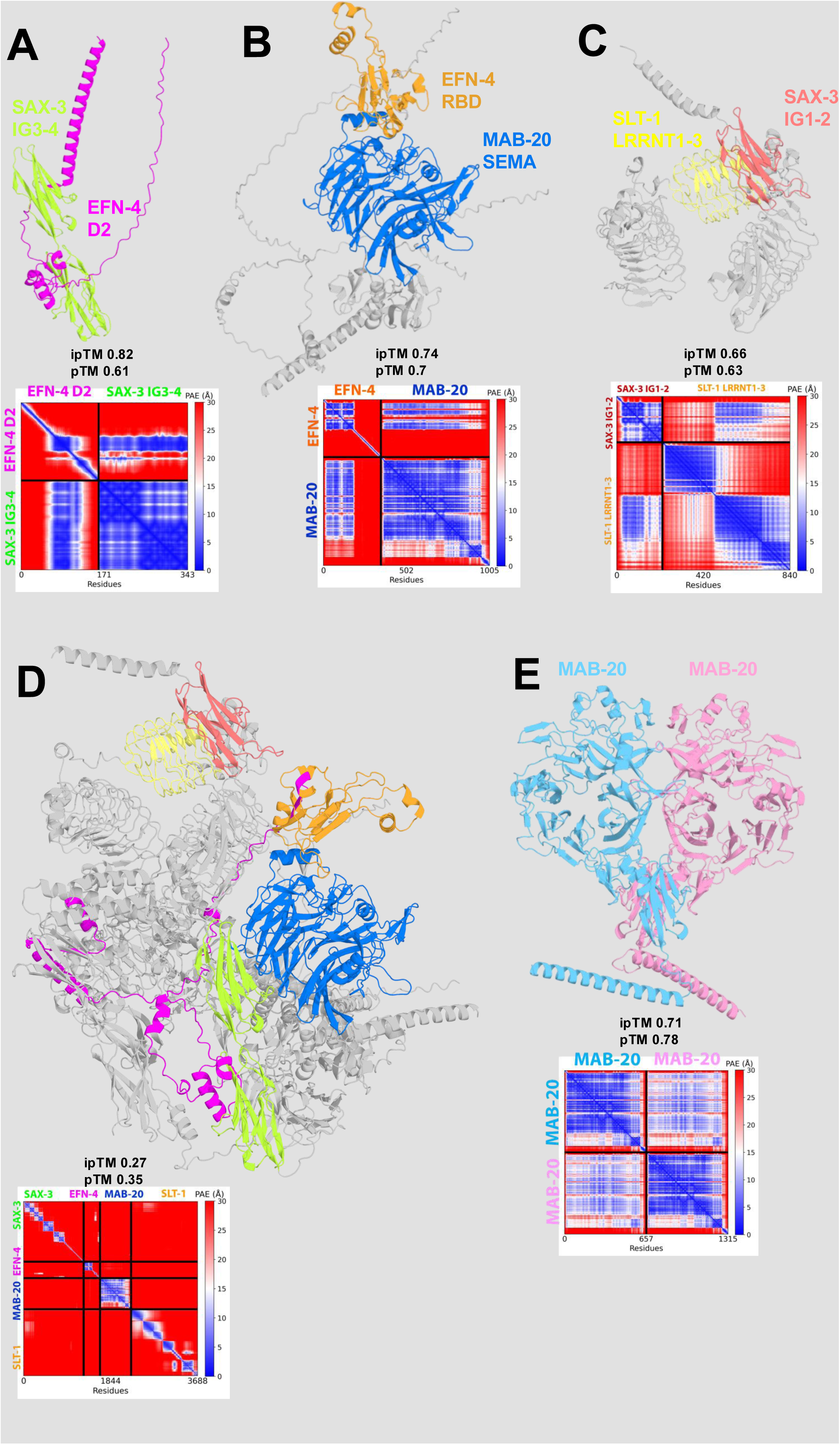
AlphaFold3 modeling of potential interactions between molecules. A) The SAX-3 IG3-4 domains interfaced with the EFN-4 D2 domain. B) The MAB-20 Sema domain interfaced with the EFN-4 RBD domain. C) The SAX-3 IG1-2 domains interfaced with the SLT-1 LRRNT 1-3 leucine-rich-repeat domains. D) A potential multimer containing the interactions described in A-C E) A MAB-20 dimer.

### *slt-1/Slit* is involved in the final QL.ap migration

The canonical SAX-3/Robo ligand SLT-1/Slit (HAO *et al*. 2001) is not significantly expressed in the QL lineage (Figure 3) (PAOLILLO *et al*. 2024; TEIXEIRA *et al*. 2025).

Previous studies did not find any AQR or PQR migration defects in *slt-1* mutants, although *slt-1* did enhance AQR migration defects of *nfm-1* mutants (JOSEPHSON *et al*. 2017). However, the subtle paradigm 2 PQR migration defects were not appreciated at the time. Upon re-scoring six different alleles of *slt-1*, we found PQR paradigm 2 migration defects in all the alleles (Table 1; 4-20%) (Figure 2C), the strongest being in the *slt-1(eh15)* putative null allele. As *slt-1* is not expressed in the Q lineages, this could be a non-autonomous role. *slt-1* is expressed in the dorsal and ventral body wall muscle cells in the posterior near the Q cells (JOSEPHSON *et al*. 2017). Posterior PQR migration defects of *sax-3; slt-1* double mutants were not significantly different than *sax-3* alone (Table 1; 20-40% combined paradigm 1 and 2). This is consistent with SLT-1 and SAX-3 acting in the same pathway.

### *unc-34/Enabled* is involved in posterior PQR migration

UNC-34/Enabled is a known cytoskeletal effector of SAX-3 signaling (YU *et al*. 2002). UNC-34/Ena has also been shown to physically interact with the second conserved proline-rich cytoplasmic motif of SAX-3 to promote actin polymerization (YU *et al*. 2002; SHEFFIELD *et al*. 2007; FLEMING *et al*. 2010; MCCONNELL *et al*. 2016; KANNAN *et al*. 2017; SHI *et al*. 2021). *unc-34* is expressed in the QL lineage (TEIXEIRA *et al*. 2025) (Figure 3), but was not affected by *mab-5* (PAOLILLO *et al*. 2024).

*unc-34(e951* and *gm104)* (FLEMING *et al*. 2010) are putative null alleles, and displayed both strong paradigm 1 and paradigm 2 PQR posterior migration defects (61% and 50%, respectively) (Table 1 and Figure 2D)*. unc-34(lq17)*, a splice site mutation and hypomorph (SHAKIR *et al*. 2006; NORRIS *et al*. 2009), displayed only paradigm 2 defects (Table 1). No anterior PQR migration was observed.

The *unc-34(e951); sax-3(gk5367)* double mutant had PQR paradigm 1 and 2 migration defects that were not significantly stronger than the single mutants alone (44% versus 61% and 46%) (see Materials and Methods for our interpretation of double mutant analysis). In fact, *unc-34(e951)* alone was significantly stronger than the double mutant (*p* = 0.0232). These genetic results are consistent with SAX-3 and UNC-34 acting in the same pathway in PQR posterior migration.

Paradigm 1 PQR migration defects in the *unc-34(e951); slt-1(eh15)* double mutant were not significantly enhanced compared to *unc-34(e951)* (12% versus 15%). Paradigm 2 defects were increased compared to each single mutant (64% versus 46% and 20%; *p* ≤ *0.016*) but were not synergistic (*i.e.* were not significantly higher than the predicted additive effect of 57%) (see Materials and Methods for our interpretation of double mutant analysis). This is consistent with *slt-1* and *unc-34* acting in the same pathway, although they might have some independent role in the third and final stage of QL.ap posterior migration.

### Genetic interactions suggest that *efn-4, mab-20, sax-3,* and *slt-1,* might converge on a common pathway

In a previous study, we showed that *efn-4, vab-8,* and *lin-17* define pathways that control posterior QL.a and QL.ap migration downstream of MAB-5 (JAIN AND LUNDQUIST 2025). To understand how these pathways might interact with SAX-3 signaling, double mutants were analyzed. As described in Materials and Methods, the proportion of defects in double mutants is compared to the predicted additive effect of the single mutants. If the proportion of the double mutant is significantly higher than the predicted additive effect, we interpret this as a synergistic genetic interaction and that the genes act in parallel, redundant pathways. Lack of a synergistic effect is consistent with the genes acting in the same pathway or possibly in independent pathways if the double mutant defects are higher than the single mutants alone.

*sax-3; efn-4* double mutants showed no significantly enhanced paradigm 2 defects compared to single mutants (Table 1). Indeed, *sax-3(gk5367)M+; efn-4(bx80)* showed significantly fewer paradigm 2 defects compared to *efn-4(bx80)* alone (53% versus 70%; *p* < 0.0001). While we do not understand the nature of this possible suppression, these results are overall consistent with EFN-4 and SAX-3 acting in the same pathway in the final QL.ap migration. Neither *sax-3(ky123)* nor *efn-4(bx80)* alone had paradigm 1 PQR migration defects, whereas the double mutant displayed 10% (Table 1). This suggests that the long isoform of SAX-3 and EFN-4 might have redundant, parallel roles in the first two QL.a and QL.ap migrations. *sax-3(zy5); efn-4(bx80)* double mutants also displayed paradigm 1 defects not observed in either single alone (Table 1).

*efn-4* was expressed predominantly in the QL.a and QL.ap lineages (Figure 3). *mab-20* was expressed in both QL.a and QL.p lineages (Figure 3). *mab-20* double mutants with *sax-3* were lethal except for *mab-20(ev574); sax-3(zy5)*, which were sterile. Therefore, *sax-3(zy5)* was maintained in this strain as a heterozygote over the *tmC30* balancer chromosome, and homozygous *sax-3(zy5); mab-20(ev574)* with *wild-type sax-3(+)* maternal contribution (*M+)* were scored. *sax-3(zy5)M+; mab-20(ev574)* double mutants displayed paradigm 1 PQR migration defects that were not significantly stronger than *sax-3(zy5)* (11% versus 4%), and paradigm 2 defects that were not significantly stronger than *mab-20(ev574)* (51% compared to 50%) (Table 1). These data are consistent with *efn-4, mab-20,* and *sax-3* acting in the same pathway in the final QL.ap migration, with the caveat of maternal *sax-3(+)* activity in *sax-3(zy5); mab-20(ev574)* doubles.

*slt-1(eh15); efn-4(bx80)* were sterile, so *slt-1(eh15)* was maintained as a heterozygous balanced line. Paradigm 2 defects of *slt-1(eh15)M+; efn-4(bx80)* double mutants did not differ significantly from *efn-4(bx80)* alone (Table 1), suggesting that SLT-1 and EFN-4 act in a common pathway. Likewise, *unc-34(e951); efn-4(bx80)* double mutants did not have significantly increased paradigm 1 or 2 defects compared to the single mutants alone (Table 1). *mab-20(ev574); slt-1(eh15)* doubles were also sterile. Paradigm 2 defects of *mab-20(ev574)M+; slt-1(eh15)* were not significantly enhanced compared to the predicted additive effect.

In sum, these genetic interactions (a lack of strong phenotypic synergy) between *efn-4, mab-20, sax-3, slt-1,* and *unc-34* are consistent with the idea that these genes act in a common pathway in posterior QL.a and QL.ap migration, with the exception being potential parallel roles of the long isoform of SAX-3 and EFN-4 in the first two stages of QL.a and QL.ap migration.

### *vab-8* and *sax-3* genetic interactions

*vab-8* was expressed predominantly in the QLa and QL.ap with less expression in QL.p (Figure 3). *sax-3(gk5367); vab-8(ev411)* double mutants displayed significantly more paradigm 2 defects and total PQR migration defects compared to the predicted additive effect (Table 1). Similarly, *sax-3(zy5); vab-8(ev411)* and *sax-3(ky123); vab-8(ev411)* showed significantly synergistic effects (Table 1). These data suggest that *sax-3* and *vab-8* might have redundant, parallel roles in posterior QL.a and QL.ap migration.

### *lin-17* and *sax-3* genetic interactions

Our previous work showed that *lin-17* acts both upstream and downstream of MAB-5/Hox (JAIN AND LUNDQUIST 2025): *lin-17* is part of the Wnt signaling pathway that activated *mab-5* expression and QL; and later, *lin-17* is upregulated by MAB-5 in the QL lineage to mediate QL.a and QL.ap posterior migration. Thus, *lin-17* mutants display some PQR neurons that fail to activate *mab-5* and thus migrate anteriorly (Table 1; 22%). *lin-17* mutants also display paradigm 1 and 2 posterior PQR migration defects (Table 1) and affect the second and third migrations of QL.a (JAIN AND LUNDQUIST 2025).

*lin-17* was expressed throughout the QL lineage (Figure 3). *lin-17(n671)* double mutants with *sax-3(gk5367)* null did not display significantly increased PQR migration defects compared to the predicted additive effects of the single mutants alone (Table 1). This is consistent with *sax-3* and *lin-*17 acting in a common pathway. However, *sax-3(zy5); lin-17(n671)* doubles did display significantly more paradigm 1 defects compared to the singles (37% versus 20% and 4%). We do not understand the nature of this interaction of *lin-17* with the hypomorphic *sax-3(zy5)* mutation, but it is possible that *lin-17* and *sax-3* have some redundant roles in the first and second steps of posterior QL.a migration.

Interestingly, *sax-3(ky123)* suppressed the anterior PQR migration defects of *lin-17* (0% versus 22%; Table 1). The nature of this interaction is not understood, but it suggests that the *sax-3* long isoform might be required for anterior migration in some contexts, or might normally prevent inhibit activation of *mab-5* in QL revealed in the *lin-17* background.

### Computational prediction of a multi-component interaction between EFN-4, SAX-3, SLT-1 and MAB-20

The Extracellular Interactome Assay (ECIA) was developed to determine the pair-wise physical interactions of the ectodomains of many secreted and transmembrane proteins in *C. elegans* (NAWROCKA *et al*. 2024), including EFN-4, MAB-20, SAX-3, and SLT-1, which are all included in the same “connected community” of physical interactors. The D2 domain of EFN-4 interacts with the first four extracellular Immunoglobulin domains of SAX-3; MAB-20 physically interacts with EFN-4; and SAX-3 interacts with SLT-1.

Using the ECIA results from (NAWROCKA *et al*. 2024) as guidance, AlphaFold3 was used to model these potential interfacing domains. The D2 domain of EFN-4 was modeled with various combinations of SAX-3 Ig domains. Ig domains 3 and 4 were found to form the strongest interface, with an Interface Predicted Template Modeling (ipTM) score of 0.82 and a Predicted Template Modeling (pTM) score of 0.61 (Figure 5A). The RBD domain of EFN-4 and Sema domain of MAB-20 were found to interface strongly with an ipTM score of 0.74 and a pTM score of 0.7 (Figure 5B). For SAX-3 and SLT-1, the strongest potential interaction was between the SAX-3 Ig1 domain and the LRRNT2 domain of SLT-1, with an ipTM score of 0.66 and a pTM score of 0.63 (Figure 5C). This was achieved by folding the Ig1-2 domains of SAX-3 with the LRRNT1-3 domains of SLT-1. In the course of modeling interactions between these molecules, we found a strong predicted dimerized fold of two copies of MAB-20, which exhibited an ipTM score of 0.71 and a pTM score of 0.78 (Figure 5E).

**Figure 5.**
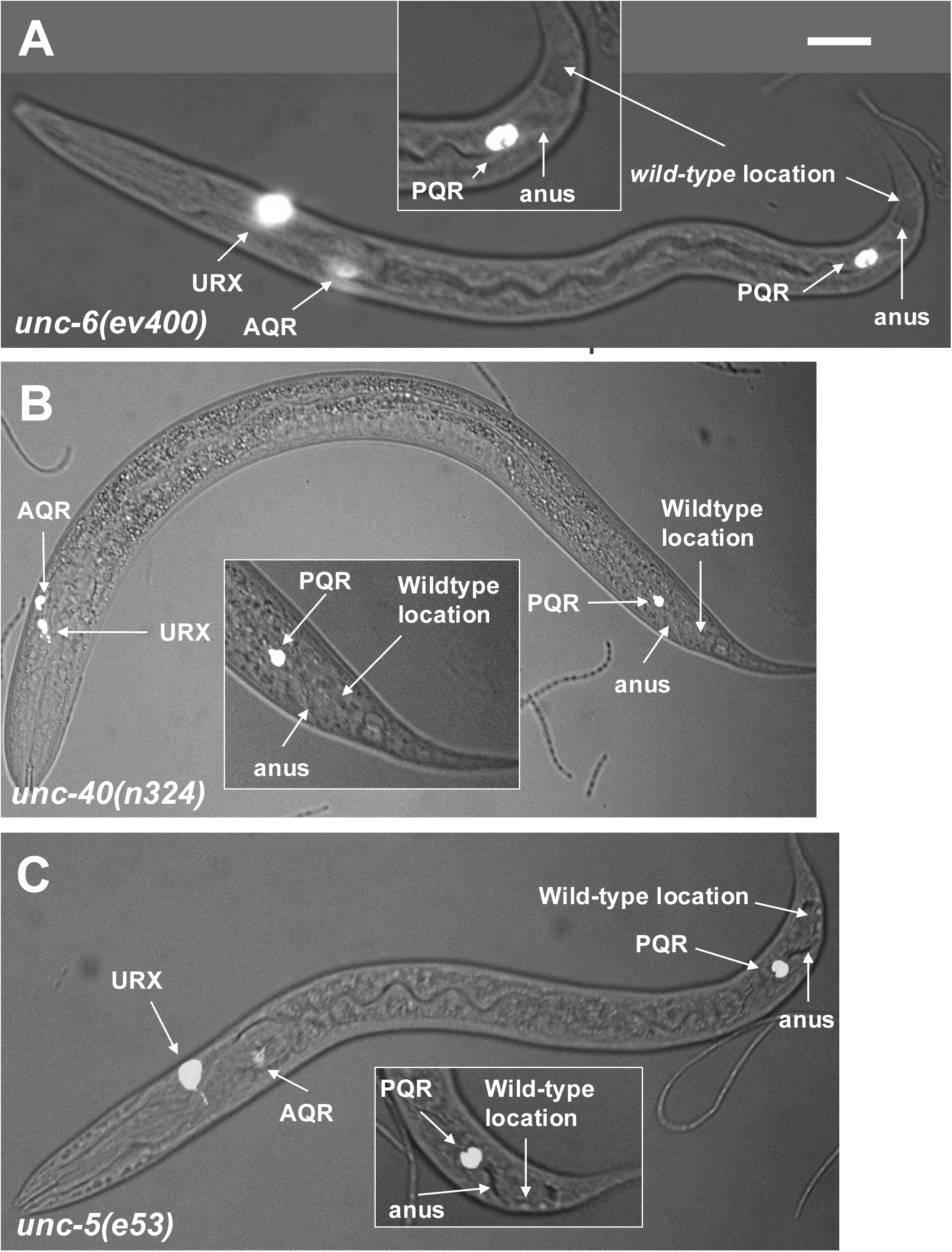
PQR migration defects in *unc-5, unc-6,* and *unc-40* mutants. Fluorescence and differential interference contrast micrographs of L4 larval stage animals with *Pgcy-23::cfp* expression in AQR, PQR, and URX left and right as described in Figure 1. The scale bar in A represents 10µm. A) An *unc-6(ev400)* animal with PQR immediately anterior to the anus. B) An *unc-40(n324)* mutant with PQR immediately anterior to the anus. C) An *unc-5(e53)* animal with PQR immediately anterior to the anus.

Finally, we used AlphaFold3 to predict if these interactions could possibly exist simultaneously in a four-way multimer. Two folds out of ninety attempts exhibited all three of these predicted interactions. From these two folds, one was selected which most closely resembled the isolated domain-specific interactions (Figure 5D). This final fold yielded a modest ipTM score of 0.27 and a pTM score of 0.35. However, when folding large protein structures with multiple interfaces and vast non-interfacing regions, these scores can be considered to be reasonable indications that these select interactions could possibly occur simultaneously. The physical interaction results from (NAWROCKA *et al*. 2024) and the modeling with AlphaFold3 here suggest that these EFN-4, MAB-20, SAX-3, and SLT-1 could interact in an extracellular multimeric complex.

### Mutations in *unc-6/Netrin* signaling affect posterior PQR migration

The results above and from (NAWROCKA *et al*. 2024) are consistent with the idea that EFN-4, MAB-20, SAX-3, and SLT-1 might form a multimeric extracellular complex that acts to control the final protrusion and migration of QL.ap posterior to the anus. As the paradigm 2 PQR posterior migration defect was not previously appreciated, it is unclear what other known guidance ligands, receptors, and extracellular molecules might be involved. UNC-6/Netrin signaling and SAX-3/Robo signaling are known to interact in axon guidance (YU *et al*. 2002; TANG AND WADSWORTH 2014). A role of UNC-6/Netrin in Q lineage migration has not been reported previously, although its receptor UNC-40/DCC controls the direction of the initial QL and QR protrusion and migration (HONIGBERG AND KENYON 2000; MIDDELKOOP *et al*. 2012; SUNDARARAJAN AND LUNDQUIST 2012).

*unc-6/Netrin* is not expressed in the Q lineages, but its receptors *unc-40/DCC* and *unc-5* are (Figure 3). *unc-6* mutants displayed paradigm 1 and 2 posterior PQR migration defects, but none migrated anteriorly (Table 1 and Figure 6A). The *unc-6(ev400)* null mutant displayed 27% combined paradigm 1 and 2 defects. The *unc-6(e78)* mutant, which more strongly affects dorsal than ventral axon guidance, displayed 50%, significantly more than *unc-6(ev400)* (*p* = 0.0013). These data indicate that UNC-6/Netrin is involved in posterior QL.a migration, and that *unc-6(e78)* might not be a simple loss-of-function.

Alone, *unc-40* mutants have anterior migration of PQR as previously described (Table 1; 57% for *unc-40(n324)* and 22% for *unc-40(e1430*) (SUNDARARAJAN AND LUNDQUIST 2012). Both mutations introduce premature stop codons (COLON-RAMOS *et al*. 2007), but *unc-40(e1430)* was significantly weaker (*p <* 0.0001). Of the PQR that did not migrate to the anterior, paradigm 1 and 2 posterior PQR migration defects were also observed (Table 1; 17% and 23%) (Figure 6B). This indicates that UNC-40 is required for posterior QL.a, in addition to its earlier role in initial direction of Q protrusion (SUNDARARAJAN AND LUNDQUIST 2012). *unc-5(e53)* null mutants displayed 26% paradigm 1 and 2 defects, and hypomorphic *unc-5(ev480)* mutants displayed only paradigm 2 defects at 9% (Table 1), significantly weaker than *unc-5(e53)* (*p* < 0.0001). Neither *unc-6; unc-40* nor *unc-5; unc-40* double mutants displayed significantly enhanced defects (Table 1), suggesting that they might act in the same pathway. These data indicate that UNC-6/Netrin and its receptors UNC-40/DCC and UNC-5 are involved in posterior QL.a migration.

### Genetic interactions between signaling pathways in posterior PQR migration

*unc-6(ev400); sax-3(zy5)* were sterile and were maintained as *sax-3* heterozygotes. *sax-3(zy5)M+; unc-6(ev400)* double mutants did not differ significantly from the predicted additive effect (Table 1). However, they did show anterior migration of PQR not seen in either single alone. This suggests that *unc-6* and *sax-3* might have overlapping roles in the initial direction of Q protrusion and migration, similar to *unc-40.* No significant difference compared to the predicted additive effect was observed in *slt-1(eh15); unc-6(ev400)* double mutants (Table 1).

*unc-40(n324); slt-1(eh15)* double mutants displayed significantly enhanced anterior migration of PQR compared to *unc-40* alone (Table 1; 92% versus 57%; *p* < 0.0001), with only 8% migrating posteriorly. Therefore, we scored only PQR neurons that migrated posteriorly or failed to migrate. Combined paradigm 1 and 2 defects of 67% were observed, significantly stronger than the predicted additive interaction (*p* < 0.0001). This suggests that UNC-40 and SLT-1 have redundant, overlapping roles in posterior QL.a and QL.ap migration. Of the cells that migrated posteriorly or failed to migrate, *unc-40(n324); sax-3(zy5)M+* also displayed significantly enhanced posterior PQR paradigm 1 migration defects compared to the predicted additive effect (Table 1).

These results suggest that *unc-40*, but not *unc-6,* acts redundantly with *sax-3* and *slt-1* in posterior QL.a migration. Furthermore, UNC-6 and SAX-3, and SLT-1 and UNC-40 might also have redundant roles in earlier initial protrusion and migration of QL, as seen in enhanced anterior PQR migration.

*efn-4(bx80); unc-6(ev400)M+* mutants showed significantly enhanced paradigm 1 posterior PQR migration defects compared to *unc-6(ev400)* alone (Table 1). This suggests that *efn-4* might have a redundant role with *unc-6* in the first two phases of QL.a and QL.ap migration. Additionally, 12% of PQRs migrated anteriorly, suggesting that *efn-4* might also act redundantly with *unc-6* in initial Q migration. Consistent with this, *efn-4(bx80)* also significantly enhanced the anterior PQR migration defects of *unc-40(n324)* (Table 2). Posterior PQR migration in *unc-6(e78); efn-4(ev480)* was not significantly different than the predicted additive effect, but did display some PQRs that migrated to the anterior.

In sum, these results are consistent with EFN-4 having some redundant function with UNC-6 and UNC-40 in posterior QL.a and QL.ap migration. Furthermore, EFN-4 might act in parallel to UNC-6 and UNC-40 in initial QL protrusion direction, and indicated by enhanced anterior PQR migration in double mutants.

*unc-6(ev400); mab-20(ev574)* double mutants were lethal and could not be scored. However, *unc-6(e78)M+; mab-20(ev574)* mutants did not display significant enhancement of posterior PQR migration compared to the predicted additive effect (Table 1), suggesting that they act in a common pathway. They did display anterior PQR migration not observed in either single mutant alone, indicating that both *efn-4* and *mab-20* have redundant roles with *unc-6* in the direction of initial QL migration.

While some genetic redundancies were found (*e.g.* in *slt-1(eh15); unc-40(n324)*), these data suggest a previously unreported convergent role of the classical axon guidance systems involving SAX-3/Robo and UNC-6/Netrin in posterior QL.a and QL.ap migration. Genetic interactions broadly reveal that these pathways act in a similar genetic pathway with EFN-4/Ephrin and MAB-20/Semaphorin. Possibly, each of these signaling molecules converges to control the third and final step in QL.ap posterior migration.

### EFN-4/Ephrin and MAB-20/Semaphorin transgenic expression can partially compensate for loss of guidance signaling in PQR migration

EFN-4/Ephrin and MAB-20/Semaphorin have been shown to partially compensate for each other’s function cell-autonomously in the third stage of QL migration, where QL.ap makes its final migration into the WT location (JAIN AND LUNDQUIST 2025).

The *Pegl-17::efn-4(+)* and *Pegl-17::mab-20(+)* transgenes were introduced into *sax-3(gk5367), sax-3(zy5), slt-1(eh15), unc-6(ev400), unc-6(e78), unc-5(e53), unc-40(n324)* (Table 3)*. Pegl-17::efn-4(+)* was able to partially but significantly rescue the migration defects of *unc-6(e78)*, *slt-1(eh15),* and *unc-5(e53)* (Table 3)*. Pegl-17::mab-20(+)* could partially but significantly rescue *slt-1(eh15), unc-5(e53), unc-40(n324)* (Table 3). These results are consistent with these molecules acting in a common pathway to control posterior PQR migration wherein transgenic expression of one molecule can partially compensate for loss of another.

**Table 3.** PQR defects in compensation experiments.

| Genotype<sup>1</sup> | PQR position<sup>2</sup> |  |  |  |  |
| --- | --- | --- | --- | --- | --- |
| *sax-3(gk5367)* | 0 | 0 | 0 | 36 | 64 |
| *sax-3(gk5367); [efn-4(+)]* | 0 | 0 | 0 | 36 | 64 |
| *sax-3(gk5367); [mab-20(+)]* | 0 | 0 | 0 | 33 | 67 |
| *sax-3(zy5)* | 0 | 0 | 0 | 18 | 82 |
| *sax-3(zy5); [efn-4(+)]* | 0 | 0 | 0 | 14 | 86 |
| *sax-3(zy5); [mab-20(+)]* | 0 | 0 | 0 | 21 | 79 |
| *slt-1(eh15)* | 0 | 0 | 0 | 20 | 80 |
| *slt-1(eh15); [efn-4(+)]* | 0 | 0 | 0 | 7 | 93* |
| *slt-1(eh15); [mab-20(+)]* | 0 | 0 | 0 | 6 | 94* |
| *unc-6(ev400)* | 0 | 0 | 0 | 27 | 73 |
| *unc-6(ev400); [efn-4(+)]* | 0 | 0 | 0 | 18 | 82 |
| *unc-6(ev400); [mab-20(+)]* | 0 | 0 | 0 | 16 | 84 |
| *unc-6(e78)* | 0 | 0 | 0 | 50 | 50 |
| *unc-6(e78); [efn-4(+)]* | 0 | 0 | 0 | 35 | 65* |
| *unc-6(e78); [mab-20(+)]* | 0 | 0 | 0 | 29 | 71* |
| *unc-5(e53)* | 0 | 0 | 0 | 26 | 74 |
| *unc-5(e53); [efn-4(+)]* | 0 | 0 | 0 | 13 | 87* |
| *unc-5(e53); [mab-20(+)]* | 0 | 0 | 0 | 10 | 90* |
| *unc-40(n324)* | 54 | 1 | 2 | 17 | 25 |
| *unc-40(n324); [efn-4(+)]* | 53 | 0 | 3 | 20 | 24 |
| *unc-40(n324); [mab-20(+)]* | 7* | 0 | 0 | 39 | 54 |

<sup>1</sup>For each genotype, 100 animals were scored on three separate occasions for a total of 300 animals.
<sup>2</sup>Numbers are average percentages from three replicates of cells at each position rounded to the nearest whole number. Paradigm 1 and 2 data were combined.
\* $p \leq 0.05$ (see Results for description).

### Heparan Sulfate Proteoglycans (HSPGs) are required for posterior PQR migration

Previous studies showed that the HSE-5 heparan sulfate epimerase was required for initial direction of Q neuroblast protrusion and migration (SUNDARARAJAN *et al*. 2015; WANG *et al*. 2015). No single HSPG gene for which mutants were available displayed this early Q migration defect, but RNAi of UNC-52/Perlecan led to AQR and PQR directional defects (OCHS *et al*. 2022). Among HSPG mutants, only *sdn-1/Syndecan* displayed weak PQR migration defects, but AQR migration defects were observed in single and double mutants (SUNDARARAJAN *et al*. 2015). Furthermore, LON-2/Glypican, SDN-1/Syndecan, and UNC-52/Perlecan have been shown to interact genetically with UNC-6/Netrin signaling (MERZ *et al*. 2003; GYSI *et al*. 2013; BLANCHETTE *et al*. 2015) and *efn-4* (SCHWIETERMAN *et al*. 2016). As the paradigm 2 posterior PQR migration defects were previously unappreciated, and the fact that *sdn-1* affects PQR migration, we scored PQR migration in viable HSPG mutants *sdn-1/Syndecan*, *lon-2/Glypican, gpn-1/Glypican, agr-1/Agrin, unc-52/Perlecan,* and *cle-1/CollagenXVIII*.

*sdn-1* was expressed robustly in the QL lineage, and *lon-2* and *agr-1* were not (Figure 3). *cle-1* was expressed in QL.ap cell, which undergoes the final migration and differentiates into PQR. *sdn-1(zh20)* is a deletion allele of the first 5 exons of the gene and is a known null, whereas *sdn-1(ok449)* is an in-frame deletion of the second exon, which contains the HS chains and hence the gene product resulting from this mutation has no HS (DIAZ-BALZAC *et al*. 2014; BLANCHETTE *et al*. 2015; LAZARO-PENA *et al*. 2018; DEGROOT *et al*. 2023). SDN-1 has been implicated as a co-receptor for SLT-1 along with SAX-3 (RHINER *et al*. 2005).

As previously reported, *sdn-1(zh20)* and *sdn-1(ok449)* mutants displayed posterior PQR migration defects (Table 4; Figure 7A). Both paradigm 1 defects (12% and 5%) and paradigm 2 defects (24% and 41%) were observed. *lon-2(e678)* displayed 11% paradigm 2 defects, and *gpn-1* displayed weak defects (1%) (Table 4). *cle-1(ju34)* displayed 9% paradigm 2 defects, with no paradigm 1 defects. *agr-1(eg1770)* and *agr-1(eg153)* showed weak paradigm 2 defects (5% and 1%) (Table 4 and Figure 7B). The viable *unc-52(e444)* mutant displayed no AQR or PQR defects as previously reported (OCHS *et al*. 2022). The *sdn-1(zh20); cle-1(ju34)* double mutant did not display significantly enhanced defects compared to the predicted additive effect, but the *sdn-1(zh20); lon-2(e687)* showed significantly increased paradigm 2 defects (Table 4) The *sdn-1(zh20); lon-2(e687); gpn-1(ok377)* triple mutant was not significantly different from the *sdn-1(zh20); lon-2(e687)* double. These results suggest that SDN-1 and LON-2 might have parallel, overlapping roles in the third and final step of QL.ap posterior migration, and that CLE-1, GPN-1, and AGR-1 have minor roles.

**Figure 7.**
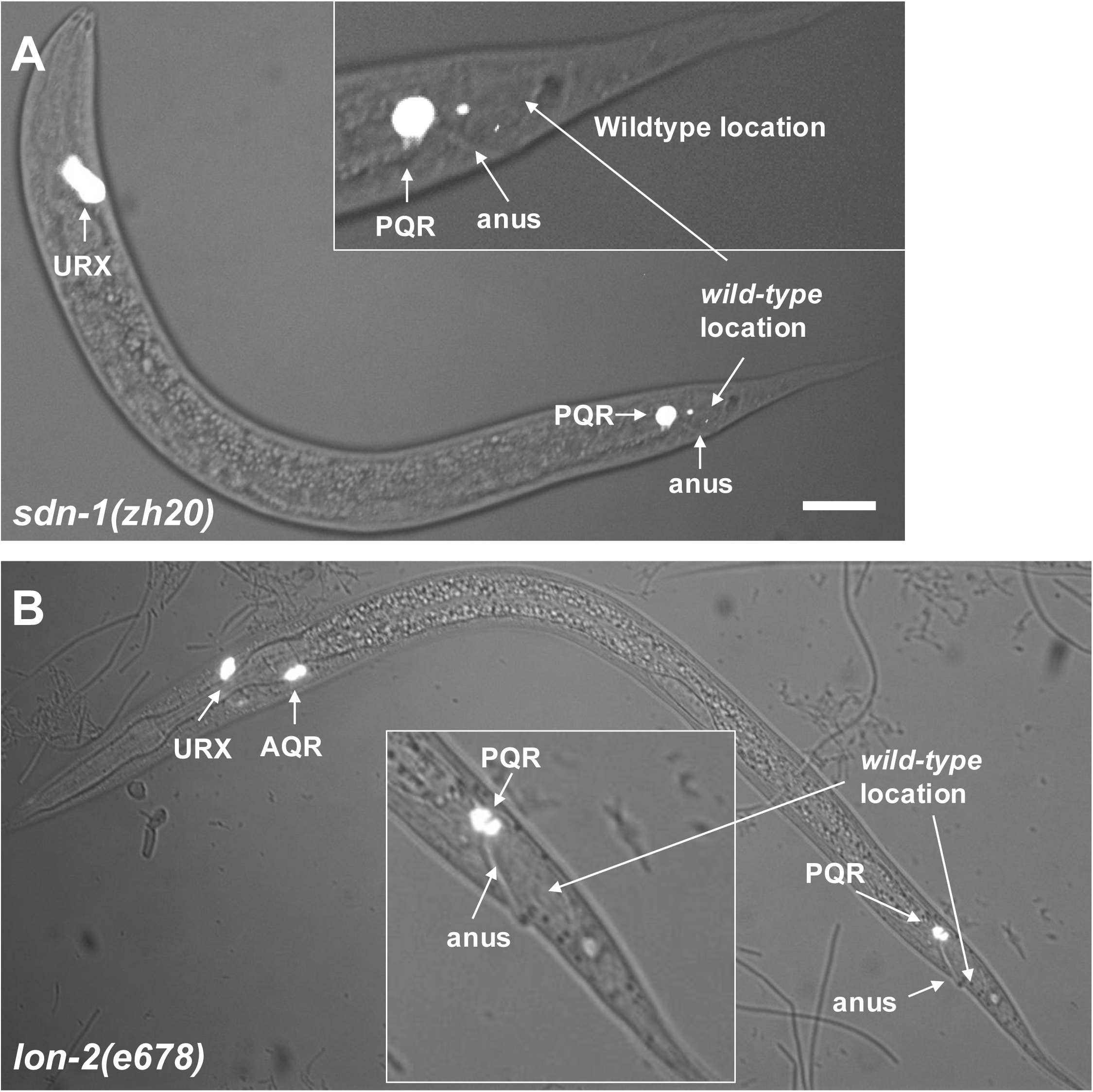
PQR migration defects in *sdn-1* and *lon-1* mutants. Fluorescence and differential interference contrast micrographs of L4 larval stage animals with *Pgcy-23::cfp* expression in AQR, PQR, and URX left and right as described in Figure 1. The scale bar in A represents 10µm. A) An *sdn-1(zh20)* animal with PQR immediately anterior to the anus. B) A *lon-1(e678)* mutant with PQR immediately anterior to the anus.

**Table 4.** PQR defects in HSPG mutants.

| Genotype<sup>1</sup> | PQR position<sup>2</sup> |  |  |  |  |  |
| --- | --- | --- | --- | --- | --- | --- |
|  | 1 | 2 | 3 | 4.1<sup>3</sup> | 4.2<sup>4</sup> | 5 |
| *efn-4(bx80)* | 0 | 0 | 1 | 0 | 70 | 29 |
| *sdn-1(zh20)* | 0 | 0 | 0 | 12 | 24 | 64 |
| *sdn-1(ok449)* | 0 | 0 | 0 | 5 | 41 | 54 |
| *lon-2(e678)* | 0 | 0 | 0 | 0 | 11 | 89 |
| *gpn-1(ok377)* | 0 | 0 | 0 | 0 | 0 | 100 |
| *cle-1(ju34)* | 0 | 0 | 0 | 0 | 9 | 91 |
| *agr-1(eg1770)* | 0 | 0 | 0 | 0 | 5 | 95 |
| *agr-1(eg153)* | 0 | 0 | 0 | 0 | 1 | 99 |
| *unc-52(e444)* | 0 | 0 | 0 | 0 | 0 | 100 |
| *sdn-1(zh20); cle-1(ju34)* | 0 | 0 | 0 | 12 | 33 | 55 |
| *sdn-1(zh20); lon-2(e678)* | 0 | 0 | 0 | 17 | 54* | 29 |
| *sdn-1(zh20); lon-2(e678);*<br>*gpn-1(ok377)* | 0 | 0 | 0 | 17 | 50 | 33 |
| *lon-2(e678); efn-4(bx80)* | 0 | 0 | 0 | 5* | 73 | 22 |
| *sdn-1(zh20)M+; efn-4(bx80)* | 0 | 0 | 3 | 21* | 70 | 6 |
| *kal-1(ok1056)* | 0 | 0 | 0 | 0 | 1 | 100 |
| *kal-1(gb503)* | 0 | 0 | 0 | 0 | 1 | 100 |

<sup>1</sup>For each genotype, 100 animals were scored on three separate occasions for a total of 300 animals.
<sup>2</sup>Numbers are average percentages from three replicates of cells at each position rounded to the nearest whole number.
<sup>3</sup>Position 4, paradigm 1 as described in Results.
<sup>4</sup>Position 4, paradigm 2 as described in Results.
\* $p \leq 0.05$ (see Results for description).

EFN-4 acts with HSPGs to control neurite outgrowth and branching, and might act in parallel to SDN-1 (SCHWIETERMAN *et al*. 2016). *sdn-1(zh20); efn-4(bx80)* showed significantly enhanced paradigm 1 defects, suggesting some redundant roles in the first and second steps of posterior QL.a migration (Table 4). *lon-2(e678); efn-4(bx80)* did not show significantly enhanced defects*. kal-1* encodes the Kallman syndrome/anosmin protein that interacts with HSPGs including SDN-1 and GPN-1 and regulates neurite formation and branching (BULOW *et al*. 2002; RUGARLI *et al*. 2002; HUDSON *et al*. 2006).

*kal-1(ok1056)* and *kal-1(gb503)* showed weak PQR migration paradigm 2 defects (Table 4), indicating a minor role in QL.ap posterior migration.

Recent *in vitro* studies have highlighted the ability of UNC-6 and UNC-5 to physically interact with heparin and facilitate the formation of large and rigid oligomeric netrin complexes that exclude UNC-40 (PRIEST *et al*. 2024). The *unc-5(wy1796 wy1996)* mutant is deficient in heparin binding due to point mutations near the first two Ig domains (PRIEST *et al*. 2024)*. unc-5(wy1796, wy1996)* displayed 14% paradigm 2 PQR migration defects (Table 1). Similar defects were seen in the *unc-5(wy1949)* mutant that affects interaction with UNC-6/Netrin (Table 1). These results suggest that UNC-5 heparin binding and interaction of UNC-5 with UNC-6 are required for the final QL.ap migration.

## Discussion

In response to MAB-5/Hox, the QL.a neuroblast and the QL.ap daughter neuron migrate to the posterior in three distinct phases, resulting in QL.ap differentiating into the PQR neuron in the phasmid ganglion posterior to the anus. Furthermore, the MAB-5-regulated genes VAB-8/KIF26, LIN-17/Fz, and EFN-4/Ephrin distinctly controlled these migrations. VAB-8 was required for each step, LIN-17 was required for steps two and three, and EFN-4 was required only for the third and final step. Failure of the third migration resulted in the previously-undescribed phenotype of the PQR neuron residing at a region immediately anterior to the anus. As this is a novel phenotype, it was not known what other genes might also control posterior PQR migration. Here we present evidence that SAX-3/Robo, UNC-6/Netrin, and heparan sulfate proteoglycans are involved in posterior PQR migration. Of all of the single and double mutants reported here, only those with *unc-40* showed AQR migration defects, which is a known role of *unc-40* (CHAPMAN *et al*. 2008). This indicates that the genetic programs described here are specific to the QL lineage and do not act in QR, although redundant roles in QR are possible.

### Many genes, each with moderate effects, control posterior PQR migration

Genetic analyses suggest that SAX-3/Robo, UNC-6/Netrin, and HSPGs might all converge on a common pathway to control the final step of QL.ap posterior migration. Each mutation alone had modest effects (∼10-30%) except for *efn-4,* which had the strongest overall effect at 70%. Furthermore, with some exceptions described below, no evidence for strong genetic redundancy was detected in double mutant analyses. *efn-4* mutants displayed the highest proportion of third-stage migration defects, and no other single or double mutant displayed significantly higher defects. High-throughput physical interaction studies showed that these molecules exist in “connected communities” that have interactions between communities (NAWROCKA *et al*. 2024). EFN-4, MAB-20, SAX-3, and SLT-1 are in one connected community, UNC-40 and UNC-6 are in another, and UNC-5 is in a third. Modeling with AlphaFold3 here supports the physical interactions between EFN-4, MAB-20, SAX-3, and SLT-1 described in (NAWROCKA *et al*. 2024).

Structural and functional studies show that Ephrins can seed the formation of distinct signaling clusters (SEIRADAKE *et al*. 2010; SEIRADAKE *et al*. 2013; NAWROCKA *et al*. 2024). Together, these data are consistent with the idea that MAB-5 drives EFN-4 expression, which serves as a “seed” for the formation of a large extracellular signaling complex that is required for the final step of posterior QL.ap migration to a position posterior to the anus. As the possible seed, EFN-4 had the strongest effect. Loss of other components led to less severe disruption, possibly because the complex can still form and have some function in the absence of any single member, except EFN-4. These data suggest a model whereby SAX-3/Robo signaling, UNC-6/Netrin signaling, and HSPGs all converge on a common pathway in response to EFN-4 to mediate the final stage of posterior QL.ap migration (Figure 8). Aside from *unc-40,* which controls initial QR protrusion and migration (HONIGBERG AND KENYON 2000; SUNDARARAJAN AND LUNDQUIST 2012), none of the genes described here caused defects in AQR migration, indicating that these genes are specifically controlling the posterior migration of PQR.

**Figure 8.**
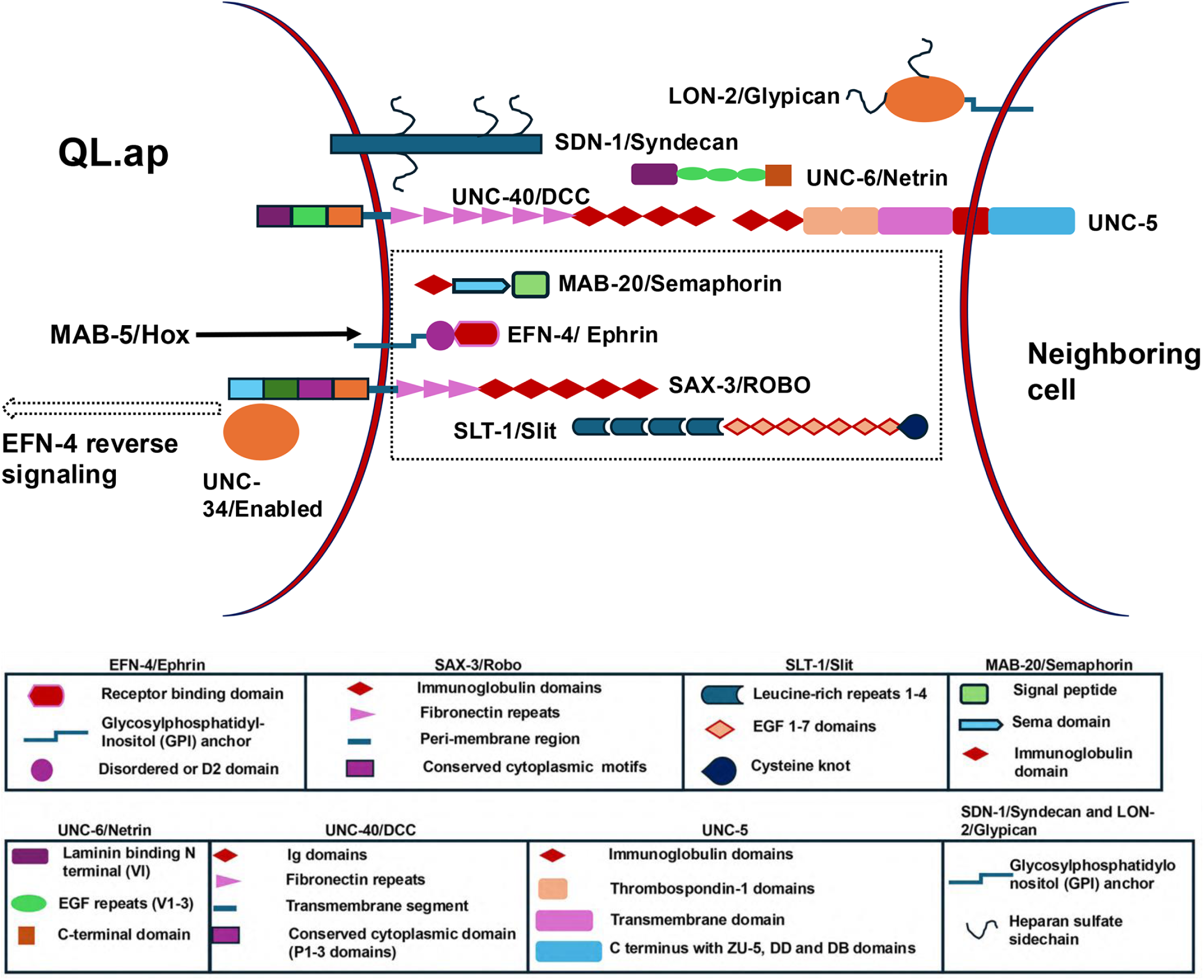
Model of EFN-4/Ephrin function downstream of MAB-5/Hox. A schematic representation of the molecules described in this work is shown, along with a legend for domains in each molecule. Molecules with the strongest effects in single and double mutants are shown. Potentially relevant molecules not shown include CLE-1/Collagen XVIII, GPN-1/Glypican, AGR-1/Agrin, and KAL-1/Anosmin. MAB-5/Hox induces EFN-4 expression in QL.ap. EFN-4 then seeds the formation a large extracellular signaling complex that, via EFN-4 reverse signaling, drives posterior lamellipodial protrusion and posterior OL.ap migration. The dashed box surrounds the molecules present in a common connected community that physically interact in the EICA interactome and might form a multimeric complex as described in Figure 5.

### Cases of genetic synergy and redundant function

Most genetic interactions described here show no genetic synergy, which is indicative of genes with redundant, parallel, overlapping functions. Redundancy is observed in several cases. First, *sax-3(ky123); efn-4(bx80)* double mutants displayed paradigm 1 PQR migration defects not observed in either single mutant alone. This suggests that while EFN-4 and the SAX-3 long isoform might act together to control the third stage of QL.ap migration, they might act redundantly to control the first and/or second stages of QL. and QL.ap migration.

Some *vab-8; sax-3* and *lin-17; sax-3* double mutants also displayed synergistic enhancement of paradigm 1 and/or 2 defects, indicating that SAX-3 might act redundantly with these molecules. VAB-8 is known to regulate cell surface localization of SAX-3, and might also be involved in the cell surface localization of an as-yet unidentified molecule that acts in parallel to SAX-3.

*unc-6(ev400); sax-3(zy5)* double mutants displayed anterior migration of PQR not seen in either single mutant alone, suggesting that SAX-3 and UNC-6 might have redundant roles in the earlier initial protrusion and migration of QL. This is the first implication of UNC-6/Netrin involvement in initial QL protrusion and migration. *sax-3(zy5); unc-40(n324)* also showed overall increase in PQR migration defects, indicating that they might act redundantly in some cases. Furthermore, *slt-1(eh15)* synergistically enhanced both anterior PQR migration of *unc-40(n324)* as well as paradigm 2 posterior PQR migration defects.

*unc-6(ev400)M+; efn-4(bx80)* and *unc-6(e78)M+; mab-20(ev574)* also displayed anterior PQR migration not seen in single mutants alone, suggesting that EFN-4 and MAB-20 also act redundantly with UNC-6/Netrin to control initial QL protrusion and migration. *efn-4(bx80)* also enhanced anterior PQR migration defects of *unc-40(n324)* suggesting that act in parallel in initial QL migration. *unc-40(n324)M+; efn-4(bx80)* also displayed synergistic defects in posterior PQR migration.

These data are the first to implicate UNC-6/Netrin, SAX-3/Robo, SLT-1/Slit, EFN-4/Ephrin, and MAB-20/Semaphorin in the initial protrusion and migration of QL, as evidenced by anterior migration of PQR in double mutants. Furthermore, UNC-40 and SAX-3 display some redundancy in posterior PQR migration. Possibly, the signaling complex can function in the absence of either receptor but is more compromised when both are perturbed. Indeed, in the ECIA conducted by (NAWROCKA *et al*. 2024), SAX-3/SLT-1 and UNC-6/UNC-40 are in distinct connected communities, with some connections between them. This might be mirrored by overlapping genetic function found here. However, no double mutant defects exceeded the level of *efn-4* alone, suggesting that EFN-4 might be required for the formation of the entire complex.

### SAX-3/Robo and SLT-1/Slit are required for posterior PQR migration

EFN-4/Ephrin belongs to the ephrin family of short-range guidance cues and resembles the A-type ephrin by topology (WANG *et al*. 1999; CHIN-SANG *et al*. 2002; MILLER AND CHIN-SANG 2012). Similar to their vertebrate counterparts, genetic evidence suggests that Ephrins in *C. elegans*, specifically *efn-1-3,* are capable of bidirectional signaling (both forward and reverse signaling) upon their interaction with the canonical Eph receptor VAB-1 (WANG *et al*. 1999; GROSSMAN *et al*. 2013; DONG *et al*. 2016). Interestingly, *efn-4* does not act with the canonical Eph receptor tyrosine kinase VAB-1 in axon guidance or embryonic morphogenesis (CHIN-SANG *et al*. 2002; DONG *et al*. 2016). VAB-1 is also not involved in posterior PQR migration as reported here.

The ECIA conducted by (NAWROCKA *et al*. 2024) found that the D2 domain of EFN-4 physically interacted with the first four Ig domains of SAX-3/Robo. Modeling with AlphaFold3 reported here suggests that Ig domains 3 and 4 form the strongest interface. Furthermore, *sax-3* mutants displayed posterior PQR defects at all three stages. These data suggest that SAX-3 and EFN-4 might act together to control the third stage, and that SAX-3 might represent the transmembrane receptor that mediates a potential autocrine EFN-4 “reverse signaling” event. Consistent with this idea, *unc-34/Enabled*, which encodes a known downstream actin cytoskeletal effector of SAX-3, also displayed posterior PQR migration defects. Recently, a novel isoform of SAX-3 lacking the Ig domains was described. The *sax-3(ky123)* mutation is predicted to only affect the Ig-domain-containing isoform, and *sax-3(ky123)* mutants displayed only defects at the third migration (paradigm 2 defects) whereas other *sax-3* mutants displayed both paradigm 1 and 2 defects. This suggests that the SAX-3 Ig domains are required for the third migration stage, consistent with interaction with EFN-4.

*slt-1* mutants also displayed paradigm 2 (third stage) PQR migration defects, suggesting that SLT-1 is also required. While SLT-1 did not interact with EFN-4 in the ECIA (NAWROCKA *et al*. 2024), it did interact with SAX-3 as expected and was in the same “connected community” with EFN-4 and MAB-20. Modeling with Aplhafold3 suggests that a higher-order complex containing these molecules is plausible, with EFN-4 interactions with SAX-3 and MAB-20, and SLT-1 interaction with SAX-3. The lack of genetic synergy between these mutants is consistent with the convergence on a common pathway, possibly a large extracellular signaling complex. As *sax-3* mutants display paradigm 1 defects, SAX-3 might be interacting with other molecules to regulate the first and second steps of posterior QL.a and QL.ap migration.

### UNC-6/Netrin, UNC-5, and UNC-40/DCC are required for posterior PQR migration

*unc-5, unc-6, and unc-40* each showed defects in posterior PQR migration, and *unc-6* and *unc-40* showed some redundancy with *sax-3, slt-1,* and *efn-4*. This is consistent with the ECIA results showing that UNC-6 and UNC-40 are in a distinct connected community of interacting molecules than EFN-4, SAX-3, SLT-1, and MAB-20 (NAWROCKA *et al*. 2024).

### UNC-6/Netrin and SLT-1/Slit might be involved in initial QL protrusion and migration

Anterior migration of PQR in *unc-6* double mutants with *sax-3, efn-4,* and *mab-20* was observed, and anterior PQR migration of *unc-40* mutants was significantly enhanced by *slt-1*. Directional defects in PQR migration are a result in the failure of initial QL protrusion and migration (CHAPMAN *et al*. 2008). Q cell migration in the wrong direction or failure to migrate could influence whether or not the cell receives the EGL-20/Wnt signal that activates MAB-5 expression and further posterior migration. These are the first data implicating secreted paracrine signaling molecules (UNC-6/Netrin and SLT-1/Slit) in the control of initial QL protrusion and migration, and the first implicating EFN-4/Ephrin and MAB-20/Semaphorin in the process. Future studies will be aimed at analyzing the defects in initial Q protrusion and migration in double mutants, as these molecules could control direction of protrusion similar to UNC-40, the ability of QL to protrude and migrate, or both.

### Heparan sulfate proteoglycans are required for posterior PQR migration

*sdn-1/syndecan* acts in parallel to *efn-4* in neurite outgrowth and branching (SCHWIETERMAN *et al*. 2016). Following the pattern of multiple genes with modest effects on PQR posterior migration, the HSPG mutants *sdn-1/Syndecan, lon-2/Glypican, cle-1/Collagen XVIII,* and *agr-1/Agrin* each had weak effects on paradigm 2 PQR migration, with *sdn-1* and *lon-2* having the strongest effect. *sdn-1* mutants also displayed paradigm 1 defects. *sdn-1(zh20); lon-2(e678)* showed synergistic paradigm 2 defects, suggesting that they might act partly in parallel. Furthermore, some potential parallel function between *efn-4* and *lon-2* and *sdn-1* in paradigm 1 defects is apparent. However, paradigm 2 defects in double mutants resembled *efn-4* alone, indicating that HSPGs might converge on a common pathway with EFN-4 in controlling the final QL.ap migration. The HSPG-interacting molecule KAL-1 had only weak effects (1%). In the ECIA interactome, KAL-1 is in the same connected community as EFN-4, SLT-1, SAX-3, and MAB-20 and physically interacts with SAX-3.

### An autocrine reverse signaling mechanism of EFN-4/Ephrin

The data presented here and in other work as described above are consistent with the model that in QL.ap, MAB-5/Hox drives the expression of EFN-4/Ephrin, which seeds the formation of a large extracellular signaling complex required for the final protrusion and migration of QL.ap past the anus into the phasmid ganglion (Figure 8). This complex likely includes factors expressed in the Q cells themselves (*e.g.* MAB-20/Semaphorin, SAX-3/Robo, UNC-40/DCC, and SDN-1/Syndecan) (Figure 3) as well as factors not expressed from the Q cells (*e.g.* SLT-1/Slit, UNC-6/Netrin, and LON-2/Glypican) (Figure 3). Interestingly, UNC-5 is not strongly expressed in QL.a and QL.ap (Figure 3). It is possible that this is a non-cell-autonomous role of UNC-5 that depends on UNC-5 expression from another cell.

*unc-5(ev480)* is a premature stop codon that affects only the long isoform containing the cytoplasmic domains (MAHADIK AND LUNDQUIST 2023), and *unc-5(ev480)* has significantly weaker PQR migration defects compared to the null *unc-5(e53)* (Table 1). This is consistent with a non-cell-autonomous role of UNC-5 wherein the cytoplasmic domains are not required. The posterior daughters QL.p and QL.pa show robust UNC-5 expression (Figure 3), which could be the UNC-5 source. Alternatively, UNC-5 could be coming from another adjacent cell, such as intestine or ventral cord neurons where it is known to be expressed (SU *et al*. 2000; DUPUY *et al*. 2007).

This EFN-4-dependent complex then signals QL.ap to extend a posterior lamellipodial protrusion and to migrate posteriorly (JAIN AND LUNDQUIST 2025). This is, in essence, reverse signaling via EFN-4/Ephrin. EFN-4 is an extracellular GPI-anchored Ephrin and has no cytoplasmic domain. The cell-autonomous transmembrane receptor components of the signaling complex such as SAX-3/Robo and UNC-40/DCC might act to transduce the signal to the cytoplasm. Indeed, UNC-34/Enabled, a known cytoplasmic cytoskeletal effector of SAX-3, is required for posterior PQR migration and *unc-34* mutants show both paradigm 1 and 2 defects, similar to *sax-3*.

EFN-4 is unlikely to be the posterior polarity of migration signal itself, as it acts in an autocrine fashion. Rather, in response to MAB-5, EFN-4 might seed the formation of a complex that drives lamellipodial protrusion and migration, with the posterior directional source of information coming from another one of the complex components. For example, *slt-1/Slit* is expressed in the anal sphincter muscle (HAO *et al*. 2001; JOSEPHSON *et al*. 2016) which is immediately posterior to QL.ap before its final migration; and *lon-2/Glypican* is expressed strongly in the posterior intestinal cells in the same region (GUMIENNY *et al*. 2007).

### Summary

This work defines a group of signaling molecules that act with EFN-4/Ephrin downstream of the MAB-5/Hox transcription factor to promote posterior PQR migration. In particular, EFN-4 is required for the third and final protrusion and migration of the QL.ap cell, which will differentiate into the PQR neuron. The genetic analysis here identified many genes and pathways involved. Single mutants in these genes have moderate effects weaker than *efn-4* mutants. This includes mutations in *sax-3/Robo* signaling, *unc-6/Netrin* signaling, and HSPGs. Some genetic redundancy was found, but a in many cases there was no significant enhancement in double mutants. This suggests that the molecules might converge on a common pathway. Our genetic results, together with the ECIA interactome study in (NAWROCKA *et al*. 2024), suggest a model by which EFN-4 seeds the formation of a large, extracellular signaling complex involving SAX-3/Robo signaling, UNC-6/Netrin signaling, and HSPGs that is required for the third and final step of posterior QL.ap migration. This could represent a case of EFN-4 reverse signaling whereby the SAX-3/Robo, UNC-40/DCC, and/or an as yet unidentified receptor transduce the signal to the cytoplasm, possibly to the actin cytoskeletal regulator UNC-34/Enabled.

## Material and Methods

### Genetics

Standard *C. elegans* techniques were used to carry out all the experiments at 20°C. Mutations used in this study were as follows: LGI: *mab-20(ev574), lin-17(n671), unc-40(n324, e1430), cle-1(ju34), kal-1(gb503, ok1056)*. LGII: *agr-1(eg1770, eg153)*, *unc-52(e444), lqIs244 [Pgcy-32::cfp].* LGIV: *efn-4(bx80), egl-20(mu39), unc-5(e53, ev480, wy1996, wy1796, wy1949).* LGV: *unc-34(e951, gm104, lq17) lqIs58[Pgcy-32::cfp].* LGX: *sax-3(gk5367, zy5, zy152, yad175)*, *slt-1(eh15, ev740, ev741), unc-6(ev400, e78), sdn-1(zh20, ok449), lon-2(e678), gpn-1(ok377)*.

### Scoring AQR and PQR locations

To score the location of both AQR and PQR neurons, we utilized the *lqIs58* and *lqIs244 [Pgcy-32::cfp* transgenes as markers, as previously described (JAIN AND LUNDQUIST 2025). Position 1 was the normal location of the AQR neuron in the anterior deirid in the head of the animal; position 2 was posterior to the pharynx; position 3 was near the vulva; position 4 was the location of the Q cell birthplace; and position 5 was the normal location of PQR behind the anus. For cells failed to migrate posteriorly and were located in position 4 two paradigms of migration defect were categorized (4.1 and 4.2) reflecting the stage at which they failed in migration (JAIN AND LUNDQUIST 2025). The first paradigm was near the QL.a birthplace or just posterior to QL.p where QL.a divides to form QL.ap, indicative of failure of the first or second migration steps. The second paradigm was PQR mispositioning immediately anterior to the anus, indicative of failure of the third and final step of QL.ap migration.

### Statistical modeling of genetic interactions

Significance of proportional difference between genotypes was determined by Fisher’s Exact text. For double mutants, the phenotypic proportion was compared to the predicted additive effect of each single mutant alone. The predicted additive effect was calculated by the formula (p1 + p2) - (p1p2), where p1 is the proportion of the first single mutant alone, and p2 is the proportion of the second single mutant alone. We interpret a significant increase as a synergistic genetic interaction, which is characteristic of genes with overlapping, redundant roles acting in parallel pathways. A non-significant difference is interpreted as an additive effect, which can be indicative of the genes acting in the same pathway or in an unrelated pathway. Each genotype was independently scored three times, 100 animals each time. The numbers shown in the tables are the percentages rounded to the nearest whole percent. For strains in which PQR migrated anteriorly, the anteriorly-migrating cells were not considered in the statistical analysis and only the PQRs that migrated posteriorly or stayed at the birth position (position 4 paradigms 1 and 2, and position 5) were considered. In *unc-40(n324); slt-1(eh15)*, only 8% of PQRs did not migrate anteriorly. Therefore, 100 PQRs that were in position 4 or 5 were scored.

### Cell-specific *sax-3* CRISPR/Cas9 genome editing

A previously described protocol for cell-specific CRISPR was utilized, wherein the expression of Cas9 was driven under a cell-specific promoters and the ubiquitous expression of sgRNA was driven under a ubiquitous small RNA U6 promoter (SHEN *et al*. 2014; OCHS *et al*. 2020). Standard ultraviolet trimethylpsoralen (UV/TMP) techniques (KAGE-NAKADAI *et al*. 2012) were used to integrate the following extrachromosomal arrays into the transgenes: unknown location *lqIs461* [*Pegl-17::Cas9/sax-3* sgRNA (25 ng/μL), *Pmyo-2::RFP* (25 ng/μL)], *lqIs462* [*Pegl-17::Cas9/sax-3* sgRNA (25 ng/μL), *Pmyo-2::RFP* (25 ng/μL)]. Two guide RNAs to *sax-3* were utilized: TCAGAGTGTAGAGTGTATCG in the third exon, and CCACCACCAACAATCGAGCA in the ninth exon.

### AlphaFold3-based prediction and modeling for protein-to-protein interactions

The EFN-4 (UniProt ID: O44516) D2 domain was folded in AlphaFold3 (VARADI *et al*. 2024) with various slices of SAX-3 (UniProt ID: G5EBF1), with the highest ipTM scoring configuration being seen between EFN-4 D2 and SAX-3 IG3-4. For the EFN-4 D2 domain, residues 168-327 were used, and for SAX-3 IG3-4, residues 227-411 were used.

Various combinations of SAX-3 and SLT-1 (UniProt ID: G5EFX6) were folded together, with the IG1-2 domains of SAX-3 interfacing best with the LRRNT1-3 domains of SLT-1, producing the highest ipTM score (seed 1575800797) from ten replicates. Residues 1-222 were used for SAX-3 and for residues 1-619 were used for SLT-1 LRRNT1-3.

MAB-20 (UniProt ID: Q95XP4) dimer was performed using seeds 1-5 using the full sequence. Seed 5 yielded the best results as reported. EFN-4 and MAB-20 full-sequences were folded using seeds 1-5, with seed 4 yielding the best results as reported.

EFN-4, MAB-20, SAX-3, and SLT-1 full-sequences were folded using seed 1-90, with seeds 42 and 63 yielding configurations reflecting three predicted interactions between EFN-4/MAB-20, EFN-4/SAX-3, and SAX-3/SLT-1

### Single-cell RNA sequencing data analysis

Q lineage single-cell RNA sequencing data publicly available at SRA (BioProject PRJNA1300790) (TEIXEIRA *et al*. 2025)were analyzed using Seurat v5.3.0 (HAO *et al*. 2024). Initial preprocessing for all three datasets was performed as described in (TEIXEIRA *et al*. 2026) without modification. Cells annotated as belonging to the QL lineage were subsetted from the merged object containing all three samples using embedded cell-type labels. Following subsetting, the data were re-normalized using NormalizeData, variable features were identified using FindVariableFeatures, and the data were scaled using ScaleData. Principal component analysis was recomputed using RunPCA. The three samples were then re-integrated using the reciprocal PCA (RPCA) method. The top seven principal components were used for graph-based clustering with the FindNeighbors and FindClusters functions. UMAP dimensionality reduction was performed using RunUMAP on the same principal components for visualization (BECHT *et al*. 2018). Gene expression was visualized using log-normalized expression values.

## Acknowledgements

This work was supported by National Institutes of Health grant R01NS115467 to E.A.L. Some strains were provided by the CGC, which is funded by NIH Office of Research Infrastructure Programs (P40 OD010440).

